# The OPAQUE1/DISCORDIA2 myosin XI is required for phragmoplast guidance during asymmetric cell division in maize

**DOI:** 10.1101/2021.08.29.458084

**Authors:** Qiong Nan, Hong Liang, Janette Mendoza, Le Liu, Amit Fulzele, Amanda Wright, Eric J Bennett, Carolyn G. Rasmussen, Michelle R Facette

## Abstract

Formative asymmetric divisions produce cells with different fates and are critical for development. We show the myosin XI protein, OPAQUE1 (O1), is necessary for asymmetric divisions during maize stomatal development. We analyzed stomatal precursor cells prior to and during asymmetric division to determine why *o1* mutants have abnormal division planes. Cell polarization and nuclear positioning occur normally in the *o1* mutant, and the future site of division is correctly specified. The defect in *o1* occurs during late cytokinesis, when the phragmoplast forms the nascent cell plate. Initial phragmoplast guidance in *o1* is correct; however, as phragmoplast expansion continues *o1* phragmoplasts become misguided. To understand how O1 contributes to phragmoplast guidance, we identified O1-interacting proteins. Maize kinesins related to the *Arabidopsis thaliana* division site markers PHRAGMOPLAST ORIENTING KINESINs (POKs), which are also required for correct phragmoplast guidance, physically interact with O1. We propose that different myosins are important at multiple steps of phragmoplast expansion, and the O1 actin motor and POK-like microtubule motors work together to ensure correct late-stage phragmoplast guidance.

## Introduction

Asymmetric divisions in plants and animals occur when a cell divides to give two daughters that differ from each other. Asymmetric divisions occur concomitantly with fate specification and are important in multicellular organisms to generate cell types of various fates, and ensure the correct relative orientations of cells required for tissue patterning. Stomatal divisions from many plant species have served as a model for asymmetric cell division and have led to the discovery of a suite of proteins required for asymmetric division and fate regulation.

Stomata are always composed of two guard cells, and may also consist of a variable number of subsidiary cells (Gray et al., 2020). Stomata from maize and other grasses are made of 4 cells: 2 inner guard cells laterally flanked by a pair of outer subsidiary cells (Nunes et al., 2020; Hepworth et al., 2018; Facette and Smith, 2012; McKown and Bergmann, 2020). Three types of divisions are required to create a stomatal complex in grasses: (1) asymmetric division of a protodermal cell to form a guard mother cell (GMC) and an interstomatal cell (Figure 1A, panel I); (2) asymmetric divisions of two subsidiary mother cells (SMCs) flanking the GMC (Figure 1A, panels III-V); and (3) the symmetric but oriented division of the GMC (not shown). Subsidiary cells are believed to contribute to the rapid stomatal movements observed in grass species (Gray et al., 2020; Nunes et al., 2020; Franks and Farquhar, 2007).

**Figure 1.**
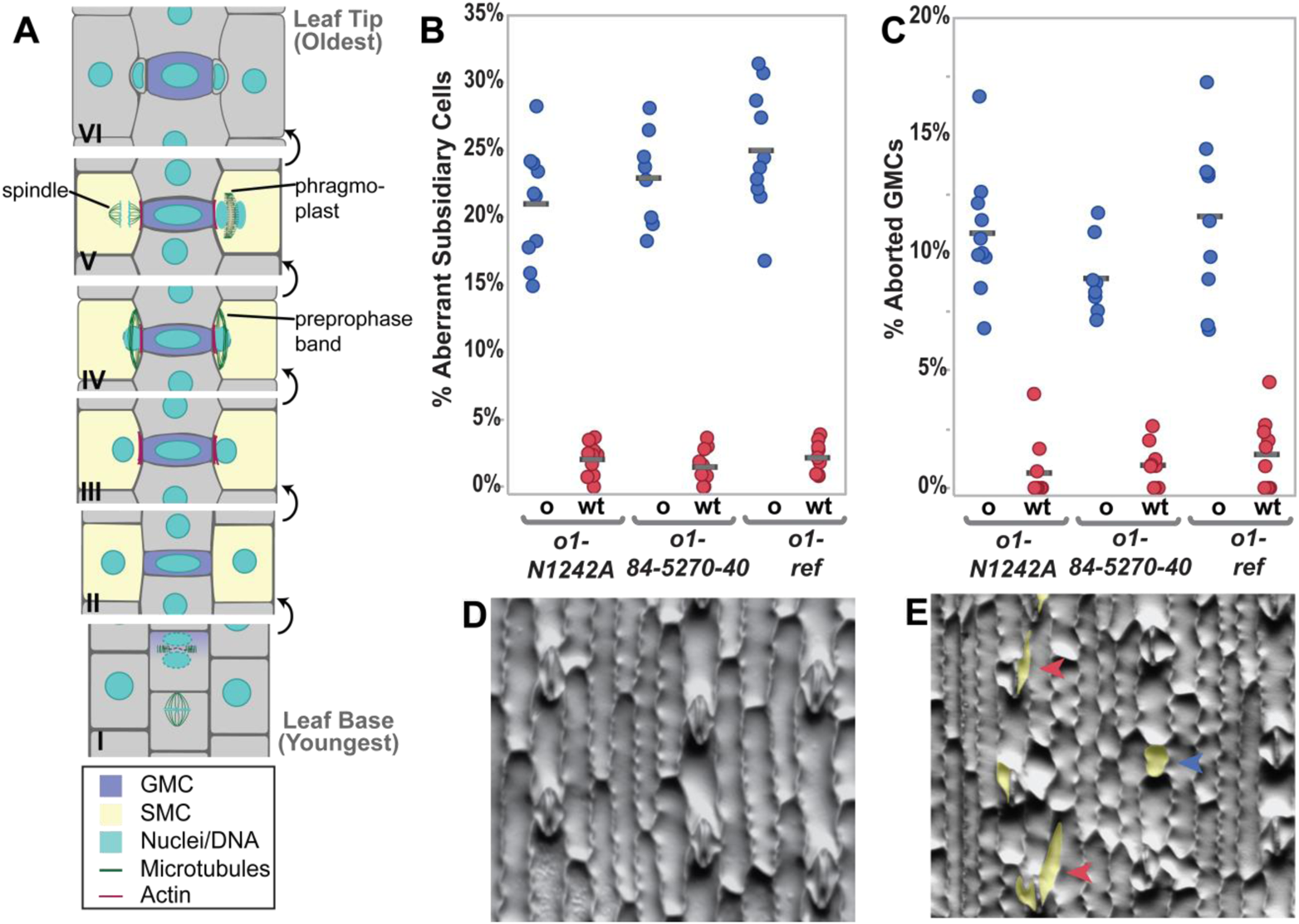
*Opaque1* is required for normal subsidiary cell formation in maize. (A) Division sequence of stomatal development in maize and other grasses. (B) Percent abnormal subsidiary cells and (C) percent aborted GMCs are increased in *o1* mutants, compared to wild type siblings. In both B & C, o = seeds that are phenotypically opaque; wt = translucent seeds. Each data point represents the percent of abnormal cells in one plant; between 100 and 200 cells were counted per plant. Grey bars indicate means. ANOVA comparing each mutant to wild type sibling yields p<0.0001 for all three alleles (D) Methacrylate impression of a wild type sibling showing normal stomatal complexes. (E) *o1-ref* mutant showing abnormal subsidiary cells (red arrowheads) and aborted GMC (blue arrowhead). Abnormal cells in E are false-colored yellow.

Stomatal development requires many coordinated cellular processes before and during the formative divisions. Mutants in grass species that fail to correctly form stomata have been identified in maize, rice, and *Brachypodium distachyon*. Grass mutants with abnormal stomatal development generally fall into two classes: mutants defective in genes important for cell fate specification, and genes encoding proteins important for physically executing the asymmetric division. Several transcription factors are important for grass stomatal divisions (Raissig et al., 2017, 2016; Wang et al., 2019; Wu et al., 2019; Liu et al., 2009). Mutants that fail to correctly execute stomatal divisions are classified based on which universal and temporally distinct phase of asymmetric division is defective: cell polarization, division plane establishment, division plane maintenance, or cytokinesis. The asymmetric division of the SMC in particular is useful as a model for dissecting the processes for asymmetric division, as they have conspicuous polarization markers and the daughter cells are morphologically distinct. Mutant genes leading to defects in cell polarization include *brick1 (brk1), brk2,* and *brk3* which encode subunits of the SCAR/WAVE complex promote actin nucleation (Frank et al., 2003; Facette et al., 2015); a pair of leucine-rich repeat receptor like proteins *pangloss2* (*pan2*) and *pan1* (Cartwright et al., 2009; Zhang et al., 2012), and ROPs (Humphries et al., 2011). Defects in these genes lead to a nuclear polarization defect and aberrant division planes. In *B. distachyon*, BdPOLAR has the opposite localization of PAN1 (it is excluded from the GMC-SMC interface) and is required for correct SMC polarization (Zhang et al., 2022). Mutants in which SMCs polarize correctly, but fail in subsequent steps include the *discordia (dcd)* mutants and *tangled1 (tan1)* (Gallagher and Smith, 1999; Smith et al., 1996; Martinez et al., 2017). *Discordia1 (Dcd1)* and its paralog *Alternative discordia1 (Add1)* encode protein phosphatase 2A subunits that are co-orthologues to the *Arabidopsis thaliana* gene *FASS/TONNEAU2,* and are required for correct division plane establishment (Wright et al., 2009; Kirik et al., 2012; Torres-Ruiz and Jurgens, 1994; Camilleri et al., 2002). Like FASS, DCD1/ADD1 are required for correct preprophase band formation. The PPB is an early division site marker, which disappears prior to metaphase. Identification of the microtubule-binding protein TANGLED1 (TAN1) answered a long-sought question of how the cortical division site was maintained throughout mitosis after the disappearance of the preprophase band (Walker et al., 2007; Smith et al., 1996). TAN1 and other division site markers are important for division plane maintenance and continuously mark the division site from prophase until telophase (Martinez et al., 2017; Cleary and Smith, 1998; Walker et al., 2007; Stöckle et al., 2016; Xu et al., 2008). The TAN1-interacting partners PHRAGMOPLAST ORIENTING KINESIN1 (POK1) and POK2 are kinesin proteins that have been characterized in *Arabidopsis thaliana,* which also mark the division site (Müller et al., 2006; Herrmann et al., 2018; Lipka et al., 2014). POK2 and TAN1 localize to the phragmoplast as well as the cortical division site (Herrmann et al., 2018; Mills et al., 2021; Bellinger et al., 2021; Buschmann and Müller, 2019).

Cortical division site markers explain how the division site is maintained throughout mitosis. However, for division plane fidelity, the phragmoplast must be correctly guided to the correct division site. The phragmoplast is composed of membranes, microtubules, actin and other accessory proteins, which starts as a disc in the center of the cell (Buschmann and Müller, 2019; Smertenko et al., 2018). Vesicles targeted to the phragmoplast fuse to form the cell plate. The phragmoplast expands in circumference until it eventually meets the existing cell wall at the cortical division site marked by TAN1, POK, and other division site markers (Smertenko et al., 2018). Phragmoplast expansion (reviewed in (Buschmann and Müller, 2019; Smertenko et al., 2018; Smertenko, 2018; Lee and Liu, 2019)) requires microtubule turnover and many different mutants affect phragmoplast stability, morphology or guidance. Phragmoplast guidance occurs in stages. The initial rapid expansion of the disc-phragmoplast is likely guided by actin networks (Molchan et al., 2002; van Oostende-Triplet et al., 2017). Phragmoplast expansion slows when one edge meets the cell cortex and eventually, through expansion, the entire circumference of the phragmoplast reaches the cell cortex (van Oostende-Triplet et al., 2017). In this later stage, microtubules at the cell cortex are incorporated into the phragmoplast, where TAN1 (and probably other division site localized proteins) are important for incorporation of cortical telophase microtubules into the phragmoplast (Bellinger et al., 2021). After phragmoplast expansion is complete, the newly formed cell plate fuses with the existing cell wall (Smertenko, 2018; Smertenko et al., 2017; Lee and Liu, 2019; van Oostende-Triplet et al., 2017). Importantly, early phragmoplast guidance, late phragmoplast guidance and phragmoplast integrity are distinct. Despite the recent progress using mutants and time lapse imaging, the precise mechanism and protein-protein interactions that promote correct phragmoplast guidance remain unclear.

Using maize stomatal precursors as a model, we wanted to understand how plant cells correctly execute asymmetric divisions. Actin-myosin networks are prominent during plant cell division generally (Sadot and Blancaflor, 2019), and specifically during the asymmetric division of the maize SMC. Actin accumulates in the preprophase band and the spindle of plant cells, and accumulates on either side of the division zone (Palevitz, 1987; Yasuda et al., 2005; Sano et al., 2005; Kojo et al., 2013; Panteris, 2008; Van Damme et al., 2007). F-actin, myosin VIII and myosin XI have been localized to the spindle and phragmoplast (Sun et al., 2018; Wu and Bezanilla, 2014; Abu-Abied et al., 2018). Actin-myosin networks play additional roles specific to asymmetric SMC divisions. In grasses and monocots with similar stomatal development, nuclear migration is driven by actin networks (Cho and Wick, 1991; Kennard and Cleary, 1997; Apostolakos et al., 2018). Actin is polarized at the GMC-SMC interface, and the SCAR/WAVE complex — which promotes actin nucleation — is required for polarization of PAN proteins (Facette et al., 2015). These observations indicate that the actin motors (i.e. myosins) are likely important for SMC divisions, and could potentially play roles during polarization, division plane establishment and/or cytokinesis. Therefore, we investigated the role of the previously identified OPAQUE ENDOSPERM1 (O1) protein in asymmetric divisions.

The *opaque* class of mutants were identified based on their seed phenotype (Neuffer et al., 1968; Gibbon and Larkins, 2005). *O1* encodes a myosin XI protein required for normal ER and protein body morphology in developing seeds, although the gene is expressed throughout the plant (Wang et al., 2012). Myosin XI family proteins are implicated in organelle trafficking and motility, cytoplasmic streaming, tip growth, auxin response, gravitropism, growth and division plane orientation (Avisar et al., 2009; Vidali et al., 2010; Madison et al., 2015; Abu-Abied et al., 2018; Talts et al., 2016; Sparkes et al., 2008; Olatunji and Kelley, 2020; Nebenführ and Dixit, 2018; Ueda et al., 2010). In *A. thaliana*, a triple myosin XI mutant has abnormal division planes, although the precise cause of these abnormal division planes is unknown (Abu-Abied et al., 2018). Notably, O1 is very similar to *A. thaliana* MYOXI-I, which is required for nuclear movement and shape (Muroyama et al., 2020; Tamura et al., 2013; Zhou et al., 2015). In *A. thaliana* stomatal precursors, pre-mitotic nuclear migration is driven by microtubule (rather than actin) networks, but post-mitotic migration of the nucleus is driven by actin networks and MYOXI-I (Muroyama et al., 2020).

We hypothesized O1 would play a role in asymmetric division of maize SMCs, perhaps during pre-mitotic polarization of the nucleus. Indeed, we found that in *o1* mutants, asymmetric divisions of both SMCs and GMCs are abnormal. However, division defects in *o1* are not a result of cell polarization defects, but rather late stage phragmoplast guidance defects. To gain insight into how O1 promotes correct phragmoplast guidance, we identified proteins that physically interact with O1. O1 interacts with maize orthologues of POK1 and POK2, in addition to actin-binding proteins and other myosins. Given their physical interaction, and the similarity of *pok* mutant phenotypes in *A. thaliana* to the phenotypes we observed in maize *o1* mutants, we hypothesize these two cytoskeletal motors work together to promote phragmoplast guidance.

## Results

### *Opaque1* is required for stomatal divisions

To determine if the OPAQUE1 myosin was involved in stomatal divisions, we examined the morphology of subsidiary cells in fully expanded juvenile leaves (leaf 4). Segregating F2 plants were phenotyped for opaque seeds and the frequency of abnormal subsidiary cells in wild type (both *o1/+* and +/+) and *o1* seedlings was calculated. Three different mutant alleles were examined. Between 20-30% of the subsidiary cells in *o1* were abnormal, while translucent seed siblings (i.e., wild type) had less than 5% abnormal subsidiary cells (Figure 1B). This is consistent with *brk, pan,* and *dcd* mutants, which also show ~25% abnormal subsidiary cells (Facette et al., 2015; Cartwright et al., 2009; Zhang et al., 2012; Gallagher and Smith, 1999). All three alleles also displayed an increase in frequency of aborted GMCs (Figure 1C). Aborted GMCs were classified when a single cell with the same morphology as a GMC was present instead of a normal 4-celled stomatal complex (blue arrow, Figure 1E). Aberrant subsidiary cells and aborted GMCs in *o1* suggest a role for this myosin in both types of stomatal formative asymmetric divisions. To confirm the defect, we compared *o1* and wild-type siblings at early stages of leaf development, when stomatal divisions were occurring. Propidium iodide (PI) was used to visualize cell walls and nuclei, and aniline blue was used to stain plasmodesmata and new cell plates. In *o1* mutants, recently formed GMCs showed abnormal division planes (Figure 2 A, B). We propose that aborted GMCs are present in expanded leaves when the GMC progenitor divides abnormally resulting in failed GMC fate specification. Abnormal division planes were also observed in recently formed SMCs (Figure 2 C, D). Importantly, no multi-nucleate cells or cell wall stubs were observed; only abnormal division planes were observed. This suggests that this specific myosin is not required for cell plate integrity, but rather it is the division plane that is affected. These data indicate a role for *O1* during asymmetric division of stomatal precursors.

**Figure 2.**
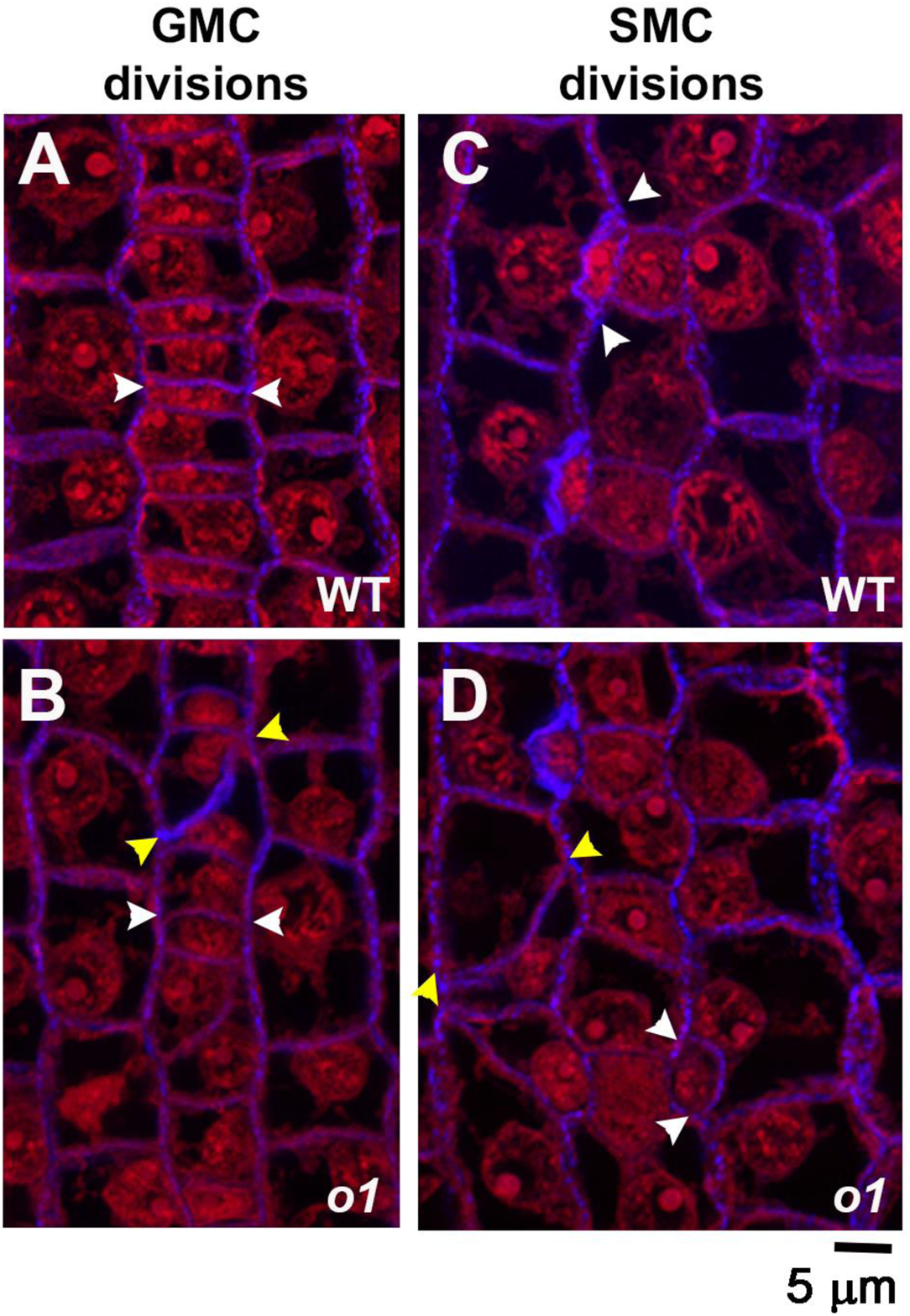
Stomatal lineage cells have abnormal division planes in *o1*. The region of developing leaf 4 undergoing stomatal divisions was dissected, fixed in FAA, and stained with propidium iodide (red) and aniline blue (blue). White arrowheads mark correct divisions; yellow arrowheads mark incorrect divisions. (A, B) Recently formed guard mother cells. (C,D) Recently formed subsidiary cells. Z-projection of 3 confocal images.

At what point does *O1* play a role in determining division plane: cell polarization, division plane establishment, division plane maintenance or cytokinesis? We examined *o1* SMCs to determine if failed polarization caused the division plane defect. Polarization in SMCs occurs in a series of ordered steps with BRK1, PAN2, PAN1 and ROP proteins each becoming polarized sequentially, and each protein is required for the next to become polarized (Cartwright et al., 2009; Zhang et al., 2012; Humphries et al., 2011; Facette et al., 2015). Actin patch formation and the nucleus are the last to polarize (Facette et al., 2015).

We assayed PAN1-YFP polarization, actin patch formation, and nuclear migration in *o1* and wild-type siblings to determine if cell polarization was normal. PAN1-YFP becomes polarized in SMCs prior to nuclear migration, and remains polarized throughout division until after the subsidiary cell is formed (Cartwright et al., 2009). We analyzed recently divided SMCs for PAN1-YFP polarization, and classified the daughter as having correct or incorrect division planes. PAN1-YFP polarized normally in all recently divided SMCs in *o1* (209/209), regardless of whether the SMC divided normally (136 cells) or abnormally (73 cells) (Figure 3, Supplemental Table 1). When a GMC progenitor cell divided abnormally and failed to form a morphologically normal GMC, adjacent cells did not polarize PAN1-YFP – presumably because GMC fate was not correctly specified (Figure 3D).

**Figure 3.**
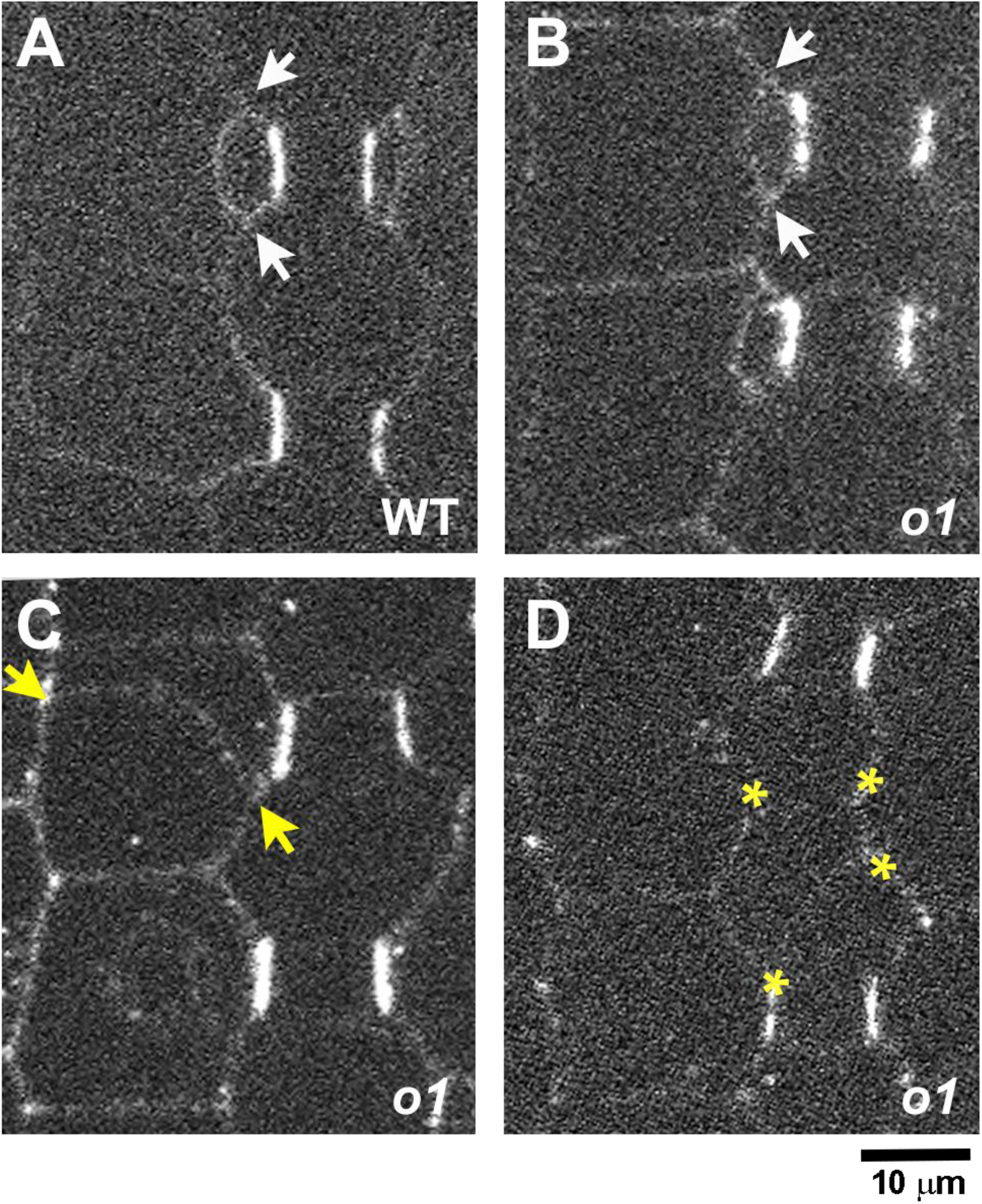
PAN1-YFP polarizes correctly in *o1*. Recently formed subsidiary cells from *o1-ref* plants and wild type siblings expressing PAN1-YFP were assayed for PAN1-YFP polarization. Arrows in A-C indicate correctly (white) or incorrectly (yellow) oriented cell walls generated from a SMC division. (A) Recently divided SMC with polarized PAN1-YFP at the GMC-subsidiary cell interface. (B) Correctly formed subsidiary cell from *o1*. PAN1-YFP is correctly polarized (C) Incorrectly oriented cell wall generated from an aberrant SMC division. PAN1-YFP is correctly polarized. (D) Incorrectly oriented cell wall generated from an aberrant division of the GMC progenitor cell. Yellow asterisks in D indicate 4 corners of a cell formed from an aberrant GMC-generating division.

We also assayed whether the polar accumulation of actin and nuclear migration to the division site occurred normally in *o1* (Figure 4, Supplemental Figure 1). We hypothesized that nuclear migration in particular may be affected in *o1* mutants, as different myosin XI isoforms were previously shown to be important for nuclear positioning in (Tamura et al., 2013; Muroyama et al., 2020; Ali et al., 2020). We assayed actin and nuclear polarization at different developmental stages. As stomatal development proceeds, GMC width increases. Therefore, GMC width can be used as a proxy for developmental state Figure 4A,C). We quantified the number of SMCs with polarized actin patches (Figure 4A,B) and polarized nuclei (Figure 4D, Supplemental Figure 1) in *o1* and wildtype siblings and found normal polarization in *o1*. Actin patch formation and nuclear polarization are the last known steps of polarization and dependent on the polarization of earlier factors, implying earlier factors that were not examined (BRK1, PAN2 and ROP) also polarize normally. Together, the data indicate that polarization is normal in *o1,* and the defect that leads to abnormal asymmetric divisions in SMCs occurs post-polarization.

**Figure 4.**
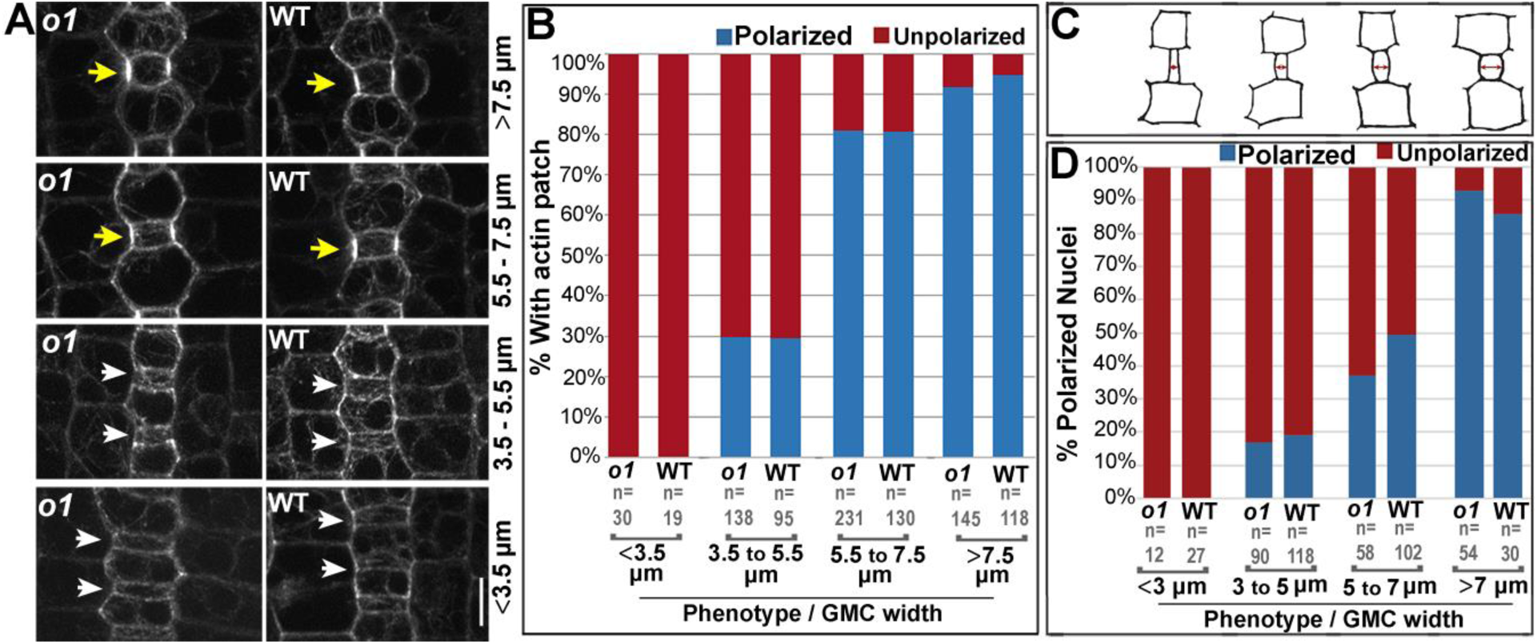
Actin patch formation and nuclear migration is normal in *o1*. (A) The stomatal division zone was examined in leaf 4 of *o1-N1242A* mutants and wild type siblings expressing the ABD2-YFP marker. GMCs in early developmental stages, found at the leaf base (lower panels) are narrow and width increases towards the leaf tip as development proceeds (upper panels). Early SMCs flanking narrow GMCs do not form an actin patch (white arrows). SMCs at later developmental stages, flanking wider GMCs, have an actin patch (yellow arrows). (B) Percent of SMCs with a polarized nucleus at progressive developmental stages in *o1-N1242A* mutants and their corresponding wild-type siblings. (C) Cartoons depicting representative cell outlines at increasing GMC widths. Red arrows indicate where GMC width was measured. (D) Percent polarized nuclei in SMCs at progressive developmental stages in *o1-N1242A* mutants and their corresponding wild-type siblings. Fisher’s exact tests comparing *o1* mutants and their respective wild-type siblings indicate no differences between mutants and wild type at each developmental stage (p>0.05 in all cases.)

### OPAQUE1 localizes to the phragmoplast

If O1 is not required for cell polarization, it must be required at a later stage of asymmetric cell division — specifically during division plane establishment, division plane maintenance and/or cytokinesis. We used immunofluorescence to detect O1 localization during cell division, to determine at what stage of the cell cycle it might be important. We generated an O1-specific peptide antibody and co-immunostained for O1 and microtubules (Figure 5 A-C; Supplemental Figure 2). Specific staining, present only in wildtype but not mutant siblings, was observed at the phragmoplast midline. O1 localized to phragmoplasts in all dividing cells we examined, including symmetrically dividing pavement cells (Figure 5A), asymmetrically dividing GMC progenitor cells (Figure 5B) and asymmetrically dividing SMCs (Figure 5C). Previously, myosin XI isoforms were shown to localize to the phragmoplast, spindle, and cell cortex (Sun et al., 2018; Abu-Abied et al., 2018), and phragmoplast localization has previously been shown for a myosin VIII protein (Wu and Bezanilla, 2014). We did not observe reproducible or specific staining in the preprophase band, spindle, or at the cortical division zone in dividing cells (Supplemental Figure 2). However, the background signal was high and therefore any faint staining (such as in the spindle midzone or cell cortex) would be difficult to detect. These data suggest that O1 plays a role during cell division, especially in the phragmoplast during cytokinesis.

**Figure 5.**
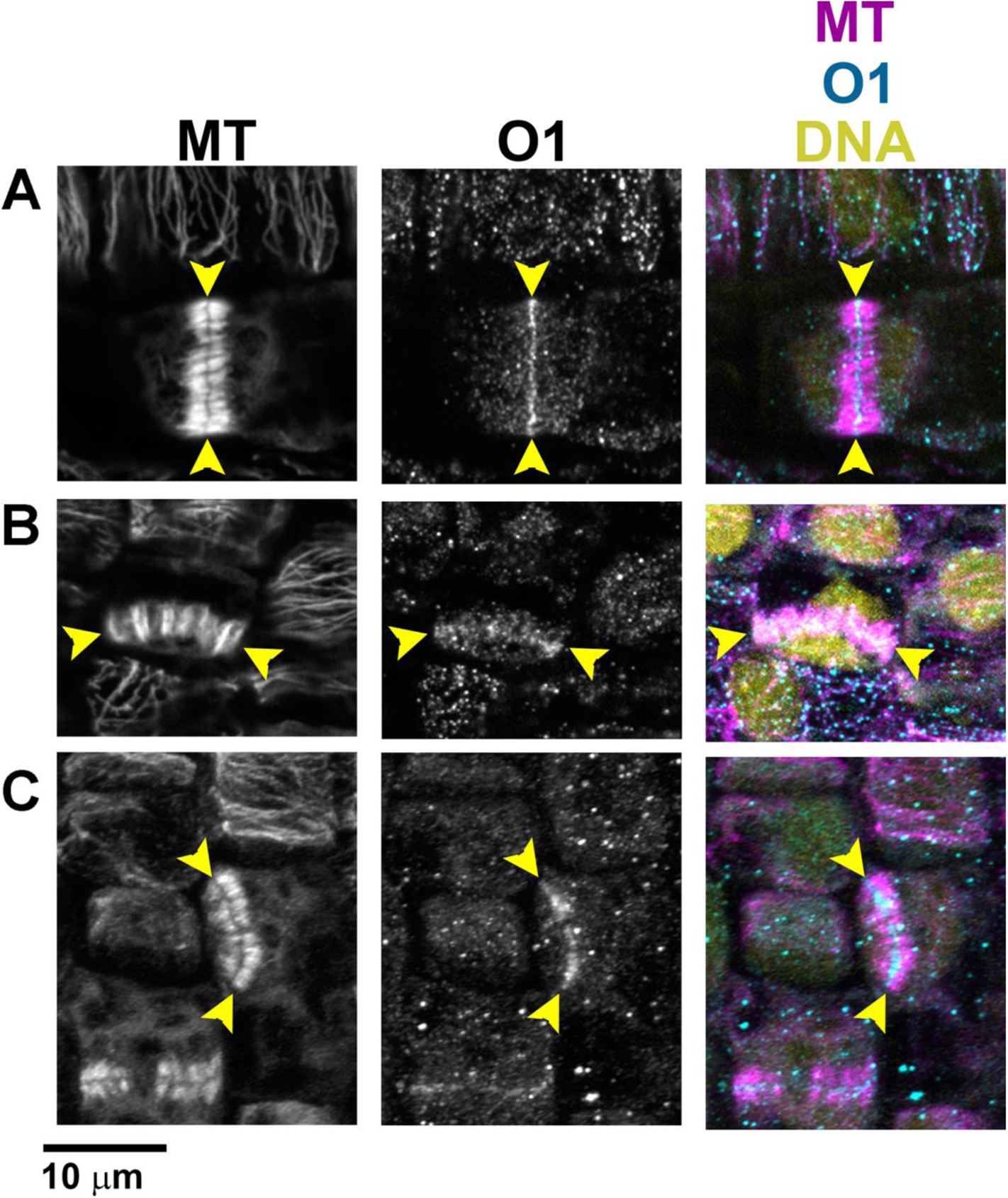
O1 localizes to phragmoplasts. Immunofluorescent detection of microtubules and O1 in wild type cells (A-C). All samples are from the division zone of developing leaf 4. O1 is detected in phragmoplasts of symmetrically dividing cells (A), asymmetrically dividing stomatal lineage cells that will form GMCs (B), and asymmetrically dividing SMCs (C). Yellow arrowheads indicate phragmoplast ends. All images were taken at the same scale.

### Phragmoplasts are misguided in *o1* mutants

Since O1 localizes to the phragmoplast and the mutant has a post-polarization defect, we wanted to know if any division structures – especially the phragmoplast – were abnormal in *o1* mutants. We examined microtubule division structures using immunofluorescence. Subsidiary mother cells from developing leaf 4 were examined (Figure 6 D-K). No abnormal preprophase bands were observed in immunostained *o1* cells (0/55) (Figure 2D,F). Spindles persist only briefly and therefore were rare, but were always normal in *o1* (0/8) (Figure 2E,G). Abnormal late-stage phragmoplasts were observed in *o1* SMCs (18/52=35%) (Figure 2 H-K). Abnormal phragmoplasts were misguided and not located at the expected site of division. However, the phragmoplast midline and microtubule alignment within the phragmoplast appeared normal and we saw no evidence for destabilized or fragmented phragmoplasts. This is consistent with our observation that no cell wall stubs were observed, and suggests normal phragmoplast assembly. The misguided phragmoplasts showed either small deviations or large deviations in the division plane. Notably, all misguided phragmoplasts appeared to be correctly oriented at one edge; i.e., one edge of the phragmoplast was always anchored at the expected SMC division site. Because O1 is an actin motor, we also examined actin in *o1* mutants. The maize ABD2-YFP marker used to assess actin patch formation does not localize to phragmoplasts (Sutimantanapi et al., 2014), therefore we used fluorescently labelled phalloidin staining on fixed cells. Phragmoplast structure and orientation in phalloidin-stained cells was similar to microtubule-stained cells – while some were normal, a subset of phragmoplasts were misguided (Supplemental Figure 3).

**Figure 6.**
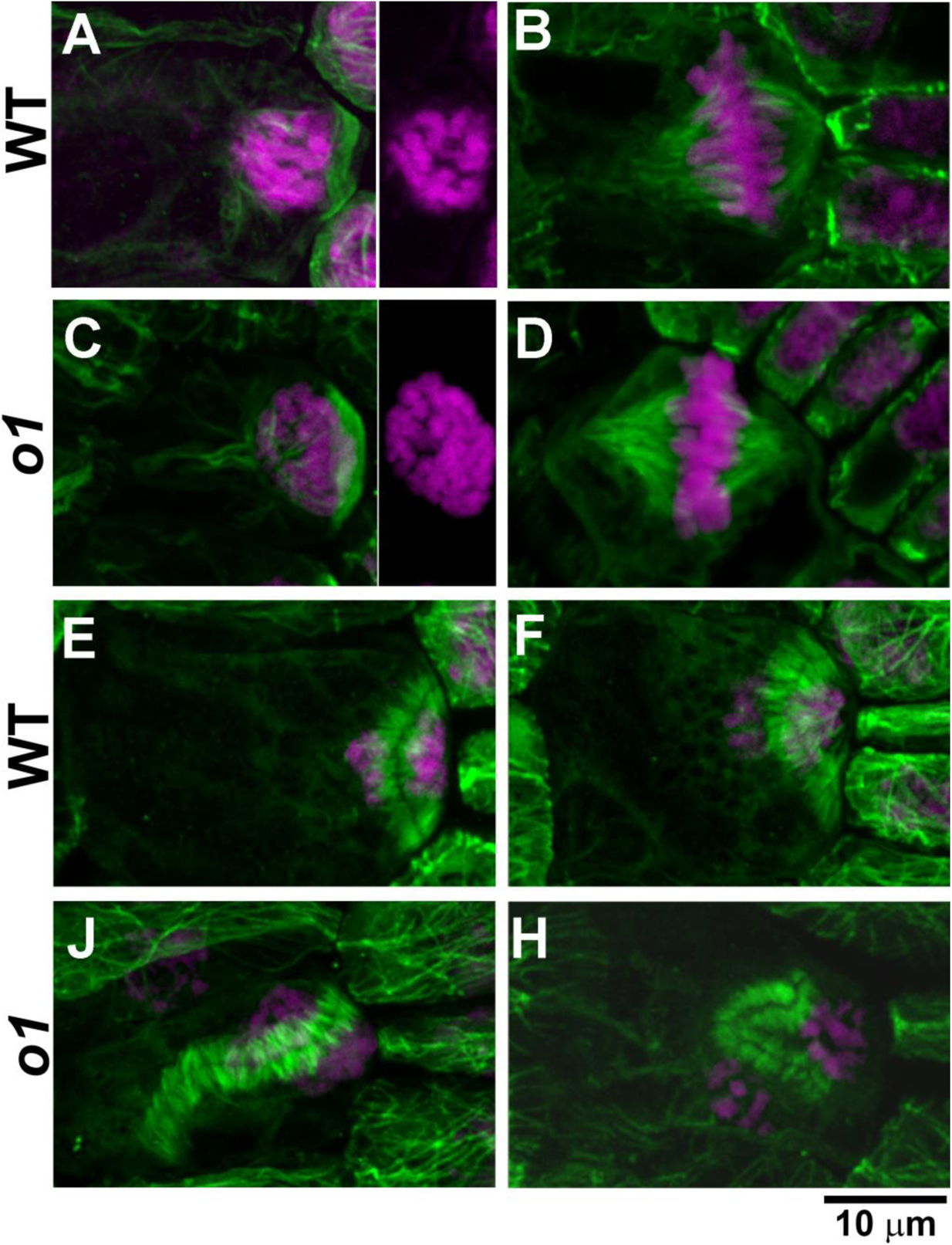
O1 is required for normal phragmoplast guidance. Immunofluorescent detection of microtubules (green) and DAP-stained nuclei (magenta) in wild type (A, B, E, F) and o1 cells (C, D, J, K). Preprophase bands in SMCs (A,C) appear similar in wild type siblings and *o1-N1242A.* Side panels in A and C show DAPI channel only to show condensed chromosomes. Spindles in SMCs (B,D) were similar in wild type and *o1* SMCs. In *o1*, SMC phragmoplasts could appear the same as wild type (E, F) or misguided (J,H). Z-projections of 40-60 images. All images shown at same scale

The *discordia* class of mutants display post-polarization defects during SMC divisions (Gallagher and Smith, 1999). Both *dcd1* and *dcd2* mutants are similar to *o1* in that they have aberrant GMC divisions, and normal nuclear polarization in SMCs (Gallagher and Smith, 1999, 2000). The gene encoding *Dcd2* had not yet been identified, but mapped to chromosome 4, and *O1* lies within the mapping interval (Supplemental Figure 4). Examination of *dcd2* seeds revealed they were opaque. Complementation crosses between *dcd2* and two *o1* mutant alleles indicate that *dcd2*, which was identified based on subsidiary cell defects (Gallagher and Smith, 1999), is allelic to *o1* (Supplemental Figure 4).

### Live cell imaging of *opaque1* mutants indicate phragmoplasts are misguided

To confirm that abnormally divided cells were a result of abnormal phragmoplast guidance, and to determine when phragmoplasts become misguided, we performed time-lapse imaging of dividing SMCs. CFP-TUBULIN (CFP-TUB) or YFP-TUB were crossed into *opaque1* (*o1*) mutants. We directly compared the location of the preprophase band, phragmoplast, and newly formed cell wall in dividing SMCs. Cells in developing leaf 5 or 6, from two alleles of *o1* plants and their corresponding wild-type siblings, were imaged from prophase until the end of cytokinesis (Figure 7, Supplemental Figure 5). In wild type, all dividing SMCs formed new cell walls that aligned with the former location of the preprophase band (n = 76 for wild type siblings) (Figure 7*A*, Movie S1). Division proceeded normally in all *o1* mutant cells until telophase. In *o1-N1242A* mutants, ~35% (n = 28/81) of divisions displayed misguided phragmoplasts (Figure 7B). In all cases, the initial site of contact between the phragmoplast edge was always aligned with the site predicted by the preprophase band. This is consistent with our e our immunofluorescence data (Figure 6), where one edge of the phragmoplast is always correctly oriented. However, as the phragmoplast continued to expand, in some cells the phragmoplast would “fall off track” and become misguided. The most severe division plane defects occurred when the phragmoplast became misguided shortly after initial contact with the existing cell wall (Movie S2); small defects occurred when the phragmoplast became misguided near the completion of expansion (Movie S3). Similar results were obtained with the *o1-5270-84* allele (Supplemental Figure 5). These data indicate that: (1) initial phragmoplast guidance, prior to first contact with the cortex, is separable from late-stage guidance and (2) in *o1,* only late-stage phragmoplast guidance is defective.

**Figure 7.**
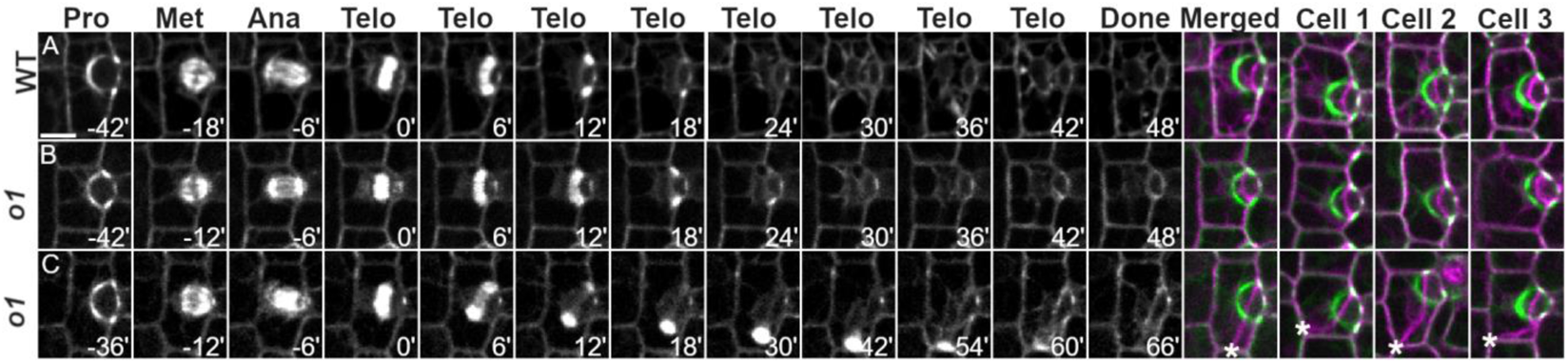
Timelapse imaging confirms a phragmoplast guidance defect in *o1*. CFP-TUB was used to observe progression of cell division SMCs from leaf 5 or 6 in *o1-N1242A* and wild type siblings. (A) Wild-type cell division. (B) Correctly oriented *o1* cell division. (C) Misoriented *o1* cell division. Pro - prophase; Met - metaphase; Ana - anaphase; Telo - telophase; Done - completed division; Merged - overlay of prophase (green) and completed division. Cell 1-3 show 3 additional representative cells. Time (minutes) listed at the bottom of each image. Misoriented cell walls are indicated by asterisks. Z-projections of 6 images. All cells displayed at the same magnification; scale bar in A = 10 µm.

### The OPAQUE1 myosin interacts with the maize orthologues of POK1/2 Kinesins

To gain insight into how O1 might be influencing phragmoplast guidance, we identified proteins that interact with O1 using co-immunoprecipitation/mass spectrometry (co-IP/MS). Membrane and membrane-associated proteins were extracted from the stomatal division zone of *o1* plants and wild-type siblings. Three biological replicates were performed using the same antibody used in immunostaining. A second set of three biological replicates was performed using a second antibody generated from an independent rabbit. Both antibodies identified 2 protein bands present in wildtype but not *o1* mutant siblings (Supplemental Figure 6). Our MS analyses isolated two isoforms of O1 that have predicted sizes corresponding to these two bands. We considered proteins enriched ≥2-fold in the wild-type samples relative to *o1* siblings, using both antibodies, to be likely O1 interactors (Supplemental Dataset 1).

High confidence interactors include many actin-associated proteins such as other myosins (including both myosin VIII and myosin XI family members), ARP2/3 proteins, and villin (Supplemental Dataset 1). Other actin-associated proteins such as fimbrin, NETWORKED, formins, or cofilin/actin depolymerizing factors (ADFs) were not identified, which suggests that the observed interactions are likely specific. The known myosin interactor MadA1 was also identified (Kurth et al., 2017).

While many actin-associated proteins were identified, the only microtubule-associated proteins identified were 2 paralogous kinesins, KIN12C and KIN12D. A third closely related kinesin, KIN12E was identified with one antibody but was just below the threshold cutoff for the second antibody, and might also be a potential interactor (Table 1). These three proteins are related to *A. thaliana* POK1, POK2 and KIN12E (Müller et al., 2006; Herrmann et al., 2018; Lipka et al., 2014; Herrmann et al., 2021). Notably, these three maize proteins were all previously identified as direct interactors of maize TAN1 (Müller et al., 2006). TAN1 (in both *A. thaliana* and maize) and AtPOK1/AtPOK2 positively mark the cortical division site during symmetric and asymmetric cell divisions and also localize to the phragmoplast (Martinez et al., 2017; Walker et al., 2007; Müller et al., 2006; Herrmann et al., 2018; Mills et al., 2022). It is possible that the interaction between O1 and maize POK orthologues is important for phragmoplast guidance.

**Table 1.** Co-IP/MS results for O1 bait protein and three related kinesin-like proteins. Co-IPs were performed using two independently generated antibodies. For each antibody, three biological replicates were done using wild type plants and *o1-N1242A* mutants as a negative control. The number of peptides identified for each replicate are separated by a slash. At least two isoforms of O1 are present in developing leaves; no peptides corresponding to O1 were identified in homozygous O1 mutants. The POK-like kinesins KIN12C and KIN12D were found only in WT samples and are considered plausible interactors. Fewer peptides were found for the related KIN12E and did not meet the threshold for a probable interactor.

| Accession | Name | Antibody 1 |  | Antibody 2 |  |
| --- | --- | --- | --- | --- | --- |
|  |  | WT Peptides | *o1* Peptides | WT Peptides | *o1* Peptides |
| Zm00001d052110_P042 | OPAQUE ENDOSPERM1 | 546/435/546 | 0/0/0 | 474/493/458 | 0/0/0 |
| Zm00001d052110_P007 | OPAQUE ENDOSPERM1 | 258/229/268 | 0/0/0 | 230/232/213 | 0/0/0 |
| Zm00001d022276_P012 | KINESIN-LIKE PROTEIN 12C | 29/3/18 | 0/0/0 | 36/56/8 | 1/0/0 |
| Zm00001d041353_P002 | KINESIN-LIKE PROTEIN 12D | 12/6/9 | 0/0/0 | 16/24/5 | 0/0/0 |
| Zm00001d034030_P017 | KINESIN-LIKE PROTEIN 12E | 3/3/3 | 0/0/0 | 7/9/6 | 4/2/0 |

Since POK proteins are division site markers, we wanted to know if the division site was being correctly maintained in *o1*. A failure to maintain the division site might explain the late-stage phragmoplast guidance defect. We used a previously characterized ZmTAN1-YFP marker line (Martinez et al., 2017) co-expressed with CFP-TUBULIN to determine if division plane maintenance was normal in *o1* mutants in different phases of mitosis. We separated telophase cells into early telophase, before the phragmoplast initially meets the cortex; and late telophase, when the defect in *o1* occurs. In wild type subsidiary mother cells, TAN1-YFP always correctly marked the predicted division plane throughout mitosis from prophase (Figure 8A) through early telophase (Figure 8B) and late telophase (Figure 8C; Supplemental Table 2; n=154 total cells). In *o1* mutants, TAN1-YFP always marked the correct division site from prophase (Figure 8D) through early telophase (Figure 8E) and late telophase (Figure 8F), even in 44 late telophase cells where the phragmoplast became misguided (Supplemental Table 2; n=231 total cells). In rare cases, TAN1-YFP was observed at an additional site during early telophase (2/231 cells; Supplemental Figure 7). Since this additional TAN1-YFP localization occurred only rarely, and we always saw correct TAN1-YFP localization in late telophase cells (when we see the phragmoplast guidance defect), ectopic TAN1 localization cannot be the primary cause of the *o1* phragmoplast guidance defect. Since the asymmetric division of GMC progenitor cells is also abnormal, we also examined TAN1 and phragmoplast localization during these divisions. Similar to SMC divisions, the division plane was correctly marked in divisions of GMC progenitor cells (Supplemental Figure 8). These data indicate that O1 is not required for correct division site maintenance and specification. Rather, O1 is required for the phragmoplast to be guided to the specified division site during cytokinesis.

**Figure 8:**
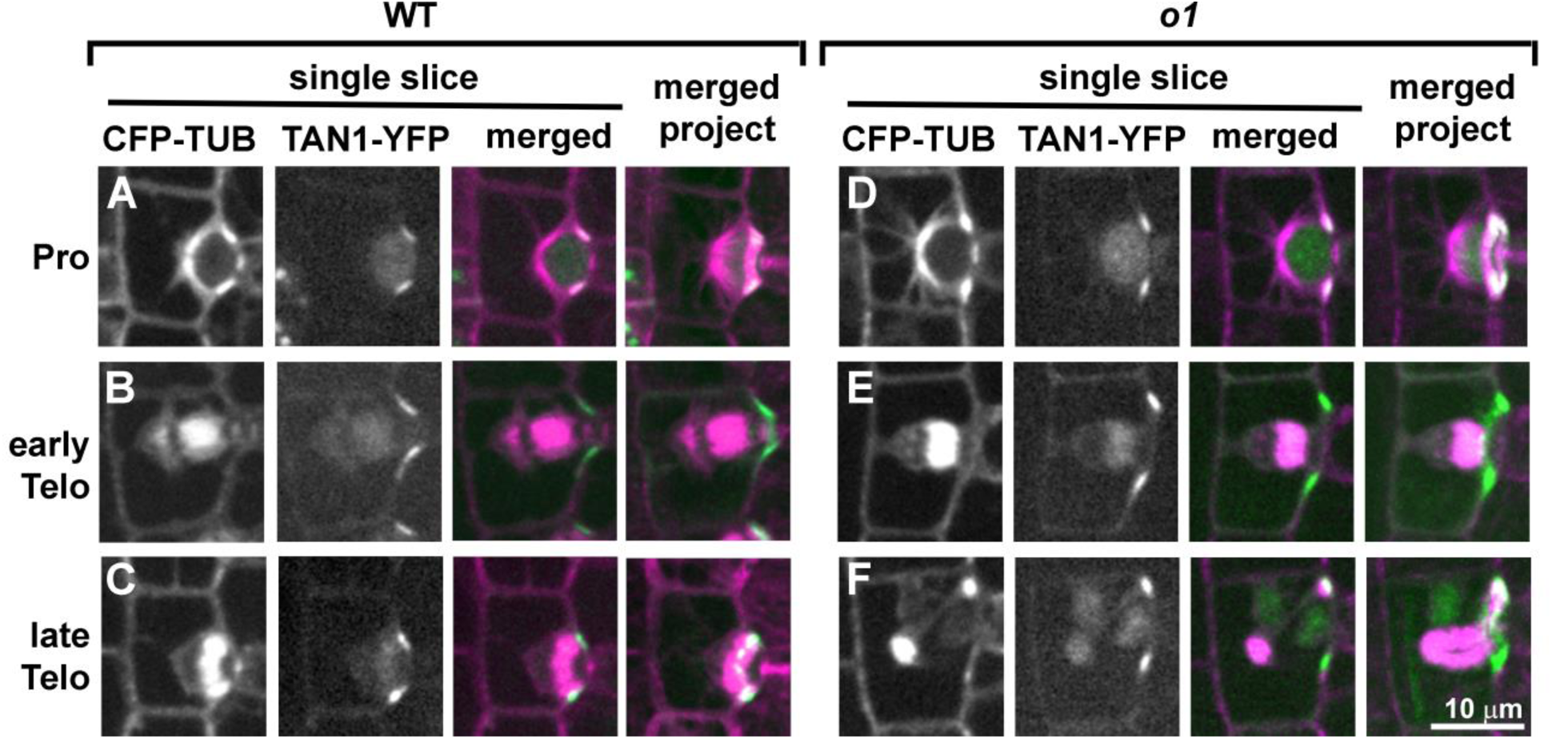
TAN1-YFP correctly marks the division plane during normal and abnormal *o1* SMC divisions. Dividing SMCs from leaf 5 or 6 in wild type siblings (A to C) or *o1-N1242A* (D-F) cells co-expressing CFP-TUB (magenta) and TAN1-YFP (green). Single planes are shown in the first three panels and a full projection is shown in the last panel. In wild-type cells, TAN1-YFP correctly marked the predicted division plane throughout mitosis including prophase (A; n=33/33), metaphase (n = 30/30), anaphase (n=12/12), early telophase (B; n=21/21) and late telophase (C; n=58/58). In *o1-N1242A* mutants, TAN1-YFP always correctly marked the division plane in prophase (D; n=85/85), metaphase (n=20/20), anaphase (n=13/13) and early telophase (E; n=32/32). In 2/32 cases, TAN1-YFP was also seen at an additional site during early telophase (see Supplemental Figure 10). During late telophase, TAN1-YFP was at the cortical division site in *o1* SMCs with correctly oriented phragmoplasts (n=38/38) and incorrectly oriented phragmoplasts (F; n=44/44). All images at same scale.

**Figure 9.**
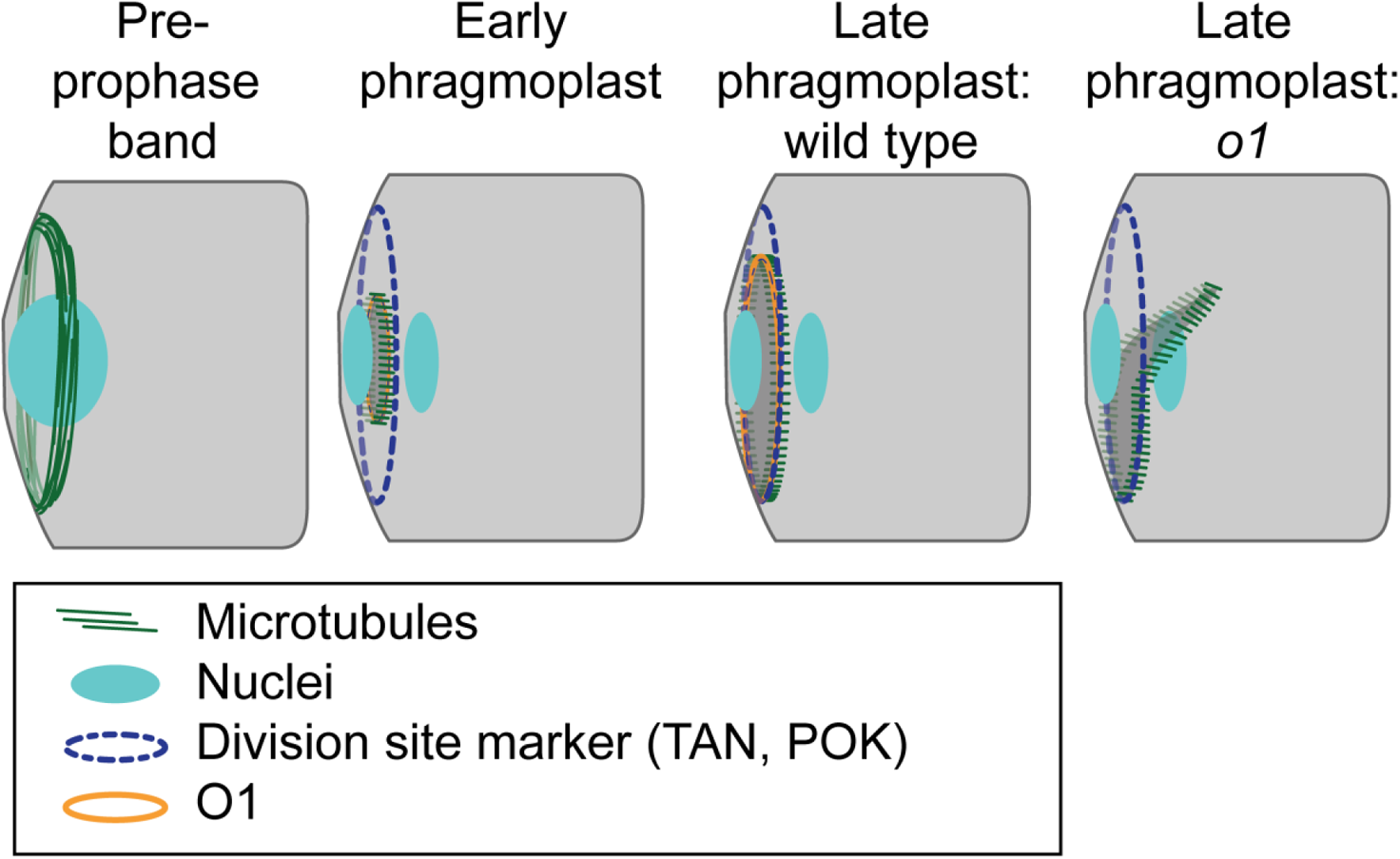
Participation of O1 in late stage phragmoplast guidance. The cortical division site is initially marked by the pre-prophase band and later by division site markers such as TAN1 and POK proteins (which are also present in the phragmoplast). In wildtype cells, the phragmoplast is guided to the division site by actin filaments and myosin VIII (Wu and Bezanilla, 2014). After meeting the cortex, the phragmoplast continues to expand and interactions are stabilized by microtubules (Bellinger et al., 2021). In wildtype, the phragmoplast fuses with the existing cell wall along the established division site, which is mediated by POK, TAN and O1. IN *o1* mutants, after initial contact the phragmoplast becomes misguided, resulting in abnormal division planes.

## Discussion

We characterized the myosin XI protein OPAQUE1/DCD2 as essential for late stage phragmoplast guidance during maize stomatal asymmetric divisions. Although myosin XIs participate in nuclear positioning in other cell types (Muroyama et al., 2020; Tamura et al., 2013), O1 is not required for pre-mitotic nuclear migration in maize SMCs - either because O1 does not play a role, or its role is masked by genetic redundancy. The asymmetric division plane defect we observed in *o1* is attributable to a late stage phragmoplast guidance defect, consistent with localization of O1 to the phragmoplast midline. This phragmoplast guidance defect is phenotypically similar to *pok1pok2* mutants (Lipka et al., 2014).

In addition to O1, many other proteins localize to the phragmoplast. Cell plate or phragmoplast localization has been observed for both myosin VIII and XI’s – some of these myosins also localize to the preprophase band, cortical division site, spindles, nuclei, and other organelles (Wu and Bezanilla, 2014; Abu-Abied et al., 2018; Zhou et al., 2015; Reisen and Hanson, 2007; Miller et al., 1995; Duan and Tominaga, 2018; Sattarzadeh et al., 2008). It was previously observed that in *A. thaliana* MYOSIN XI-K localizes to the cortical division site, spindle, and phragmoplast; in the same paper a triple myosin XI knockout led to aberrant division planes in the root (Abu-Abied et al., 2018). However, no clear mechanism was shown how XI-K might lead to aberrant division planes. Indeed, given the complex localization pattern, there are several potential roles for myosins during division. We show that in the case of O1, the division plane defect is specifically due to late stage phragmoplast guidance. Although the role for O1 appears to be specific, it is plausible that myosins have other roles in cell division and that multiple myosins might fulfil the same role. Like XI-K, a moss myosin VIII is localized to multiple mitotic structures, and mutants also has a phragmoplast guidance defect, including slowed vesicle delivery (Wu and Bezanilla, 2014). Treatment with a general myosin inhibitor alters division planes (Molchan et al., 2002), cellular polarity (Holweg et al., 2003), and even resulted in incomplete divisions (Molchan et al., 2002). We did not observe polarity defects or failed divisions; together these data suggest multiple roles for myosins during division and cytokinesis. Interestingly, we observed physical interactions between O1 and multiple other myosins, including both class VIII and XI myosins. Careful phenotypic analyses of different myosin mutants will help unravel each of the roles actin-myosin networks play during cell division.

Our data confirm that phragmoplast guidance is dynamic and that different proteins likely play different roles at different stages. Dividing cells treated with caffeine are unaffected in early-stage phragmoplast guidance but the phragmoplasts disintegrate at later stages (Valster and Hepler, 1997) and often results in multi-nucleate cells. This led to the conclusion that phragmoplast guidance occurs in (at least) two steps (Valster and Hepler, 1997), and is coupled with the observation that actin filaments connect the phragmoplast leading edge and the cell cortex (Wu and Bezanilla, 2014; Valster and Hepler, 1997). Indeed, studies of phragmoplast guidance in tobacco BY-2 cells indicate that rates of phragmoplast expansion vary, and slow considerably at last stage expansion when the leading edge first strikes the cortex (van Oostende-Triplet et al., 2017). At this final stage, the phragmoplast is more sensitive to actin depolymerization via latrunculin B (van Oostende-Triplet et al., 2017). It is also during this late stage when cortical telophase microtubules are incorporated into the phragmoplast (Mills et al., 2021). Since the phragmoplast always correctly meets the cortical division site at the initial site of contact in *o1* mutants, it is this last stage of expansion where O1 plays its role.

The identification of O1-interacting partners provides clues as to O1 functions. Since co-IP data will report both direct and indirect interactions, so some observed interactors may be indirect. O1 interacts with many proteins, including actin binding proteins, confirmed myosin binding partners and other myosins. Our data is consistent with prior observations of interactions between myosin-MadA1 (Kurth et al., 2017) myosin-(super)villin (Smith et al., 2013) and myosin-calmodulin/calmodulin like (Shen et al., 2016). We also observed interactions with O1 and two closely related Kinesin 12 proteins that are similar to AtPOK1. Even though many kinesins have been localized to the phragmoplast midline (Smertenko et al., 2018), we only identified kinesins 12D and 12E as O1-interactors. Moreover, the only other two microtubule-associated binding proteins we observed were IQ-domain proteins, recently shown in *A. thaliana* to interact with POK kinesins (Kumari et al., 2021). Mutations in *pok1* and *pok2* lead to misguided phragmoplasts similar to those observed in *o1* (Herrmann et al., 2018; Lipka et al., 2014). Mutations in other class 12 Kinesins, such as *A. thaliana* PAKRP1 and PAKRP2 (Kinesins 12A and 12B) and *P. patens* KINID1a and KINID1b (Kinesins 12B) lead to severe phragmoplast structural defects not seen in *pok* or *o1*, despite similar localization to the phragmoplast (Lee et al., 2007; Pan et al., 2004; Hiwatashi et al., 2008). Indeed, mutations in several phragmoplast-localized proteins result in defects that alter phragmoplast structure, often leading to cell wall stubs and multinucleate cells (Bannigan et al., 2007; Ho et al., 2011; Schmidt and Smertenko, 2019; Zhang et al., 2018, 3; Müller et al., 2004). The similar phenotypes of *pok* and *o1* coupled with their physical interaction suggests they work together to ensure correct phragmoplast guidance.

How might O1 and the POK-like kinesins work together to ensure proper phragmoplast guidance and disassembly? During late and slow phragmoplast expansion (van Oostende-Triplet et al., 2017), the phragmoplast falls “off-track” and becomes misguided in *o1*. Since POK proteins mark the cortical division site, and O1 localizes to phragmoplast midline, a potential model is that physical interactions between POKs at the division site and O1 at the phragmoplast midline help the phragmoplast “zipper-up’ around the cell cortex at the time of cell plate fusion. O1 and POK proteins may mediate interactions between the actin and microtubule cytoskeletons, to promote fusion of the phragmoplast at the cell wall. An alternative (or additional) model is that POK-like kinesins interact with O1 within the phragmoplast to coordinate microtubule and actin functions that promote slow phragmoplast expansion or phragmoplast disassembly once it reaches the cell cortex.

## Materials and Methods

### Plant material and growth conditions

The accession number for *O1* is Zm00001d052110 (B73 RefGen_v4, AGPv4) or Zm00001eb193160 (B73 RefGen_v5). Three *o1* mutant alleles (*o1-ref, o1-N1242A* and *o1-84-5270-40)* used for this study were obtained from the Maize Genetics Cooperation stock center. The accession number for *O1* is Zm00001d052110 (B73 RefGen_v4, AGPv4) or Zm00001eb193160 (B73 RefGen_v5). Mutant alleles were backcrossed into B73 inbred wild-type one to four times, and then selfed. In all experiments, segregating *o1* mutants were analyzed and compared to their corresponding wild-type siblings (grown side-by-side) as controls. Segregating plants were classified by their seed phenotype.

Mutant *o1* plants were crossed with various fluorescent protein-tagged maize lines generated by the Maize Cell Genomics Project (described at http://maize.jcvi.org/cellgenomics/index.php). PAN1-YFP (Humphries et al., 2011), actin maker line YFP-ABD2-YFP (Mohanty et al., 2009), TAN1-YFP (Martinez et al., 2017) or tubulin maker lines CFP-β-tubulin and YFP-α-tubulin (Mohanty et al., 2009), were crossed into *o1* homozygotes, and the F1 progeny were backcrossed with *o1* homozygotes to obtain progeny expressing the fluorescent marker and segregating homozygous mutant (*o1/o1)* and phenotypically wild-type heterozygotes (*o1*/+) were used for performing experiments.

Plants used for phenotypic analysis and image were grown for 10d to 14d in a greenhouse maintained between 72°F and 90°F with under natural light in greenhouses at the University of New Mexico, University of Massachusetts Amherst, or University of California, Riverside.

### Stomatal defects in expanded leaves

To quantify subsidiary cell and guard mother cell defects, *o1* homozygous mutants and wild-type siblings from self-pollinated *o1-ref/+*, *o1-N1243/+* and *o1-N1478A/+* were classified via their seed phenotypes. Impressions of fully expanded leaf 4 of o*1* homozygous and corresponding wild-type siblings were prepared using cyanoacrylate glue (Allsman et al., 2019) and imaged on a Nikon stereo microscope.

### Confocal Microscopy

Instruments are described here; further details on specific experimental protocols are given below.

For O1 immunostaining and nuclear polarization: Images were collected using a Zeiss LSM710 with a 63× oil immersion objective. Aniline blue was excited at 405-nm with a violet blue laser, PI was excited using the 568-nm laser line and emission filter 620/60.

For PAN-YFP and ABD2-YFP localization: ABD2-YFP-ABD2 and PAN1-YFP images were acquired with a custom spinning disk confocal microscope (3i) equipped with a Yokagawa W1 spinning disk with 50 um pinholes, iXon Life 888 EM-CCD camera (Andor) using 150 EM-CCD intensification, ASI piezo stage and solid-state lasers. YFP Images were acquired using a 60X (1.2NA) silicone immersion objective, YFP fluorescence was excited by a 514 nm 100 mW solid state laser at 6% with a dichroic excitation filter (Chroma) and a 542/27 emission filter (Semrock).

For microtubule immunostaining and phalloidin staining: Immunolocalization and actin localization experiment images were collected with a Nikon A1R with a 60X (NA 1.40) oil immersion objective. Alexa Flour 488 and Alexa Flour 568 were excited at the appropriate wavelengths of 488-nm and 568-nm, respectively, emission filters were 525/50 nm for Alexa Flour 488 and 595/50 for Alexa Flour 568.

For live imaging of CFP-TUBULIN, YFP-TUBULIN and TAN1-YFP: Time-lapse imaging was performed using a custom-built spinning disk confocal microscope (Solamere Technology) with a Yokogawa W1 spinning disk (Yokogawa), EM-CCD camera (Hamamatsu 9100c), and an Eclipse Ti-U (Nikon) inverted microscope. A 60X water immersion lens (1.2 NA) was used with perfluorocarbon immersion liquid (RIAAA-678, Cargille). The stage was controlled by Micromanager software (www.micromanager.org) with ASI Piezo (300 µm range) and 3 axis DC servo motor controller. Solid-state lasers (Obis from 40-100mW) and standard emission filters (Chroma Technology) were used. For CFP-TUBULIN, a 445 laser with emission filter 480/40 was used. For YFP-TUBULIN and TAN1-YFP, a 514 laser with emission filter 540/30 was used. All image analyses and figure preparations including cell measurements and processing was performed using ImageJ/FIJI (Schindelin et al., 2012) Adobe Photoshop CS6 or GIMP using only linear adjustments.

### Polarization measurements

To analyze PAN1 polarization, the basal 0.5-2.5 cm of leaf 4 of *o1-ref* homozygotes and heterozygous wild-type sibling plants expressing PAN1-YFP were examined. Z-stacks of the stomatal division zone were collected using a spinning-disk confocal microscope (3i) described above using a 60X silicone-oil immersion lens, YFP settings and a 500ms exposure. Recently formed subsidiary cells adjacent to a GMC that had not yet divided were assayed for polarity. PAN1-YFP polarization was scored by eye by comparing fluorescence intensity at the GMC-SMC interface and the adjacent SMC cell membrane. Cells were scored as having divided “normally” if both ends of the newly formed cell wall met the angled wall of the subsidiary cell.

To analyze F-actin polarization, the basal 0.5-2.5 cm of leaf 4 of *o1/o1* and wild-type o1/+ expressing YFP-ABD2-YFP (Mohanty et al., 2009) were examined using a spinning-disk confocal microscope (3i) described below using YFP settings and a 100ms exposure. Actin polarization was scored by eye by comparing fluorescence intensity at the GMC-SMC interface and the adjacent SMC cell membrane. GMC widths were measured using FIJI (Schindelin et al., 2012). Cell counts were then binned by GMC width and the % of cells with an actin was calculated.

To analyze nuclear polarization, double staining using aniline blue and propidium iodide of fixed tissue was performed. The stomatal division zone (basal 0.5-2.5 cm of unexpanded leaves) from leaf 4 of *o1* plants and wild-type siblings was isolated and fixed with FAA (3.7% formaldehyde, 5% acetic acid, 50% ethanol) for 1 h. Tissues were stained with 0.1% (w/v) aniline blue in PBS buffer at pH 11 for 30 min. After rinsing with PBS buffer, the tissues were stained with propidium iodide (10 µg mL^−1^ in water) and mounted on slides.

### Anti-O1 antibody generation

Custom rabbit antibodies were obtained from Pacific Immunology (Ramona, CA). Two peptides (Cys-NSEPKHIYESPTPTK and NSEPKHIYESPTPTK-Cys) were co-injected into two separate rabbits. The resulting sera were affinity purified against both peptides, according to (Cartwright et al., 2009). The two antibodies were named O1-11759 (Antibody 1) and O1-11760 (Antibody 2).

### Immunolocalization and phalloidin staining

Dual labelling of O1 and microtubules was performed as previously described, with minor modifications (Cartwright et al., 2009; Nan et al., 2019). The basal 0.5-2.5 cm of leaf 4 from *o1-N1242A* and wildtype siblings or *o1-N1242A* and wildtype siblings was used. Immunolocalization and phalloidin staining were performed separately. For immunolocalization, the dilutions used for rabbit anti-O1 (antibody O1-11759) and mouse anti-tubulin (Sigma Aldrich) antibodies were 1:1000. Alexa Flour 488-conjugated anti-mouse and Alexa Flour 568-conjugated anti-rabbit (Invitrogen) were used at a dilution of 1:500. Samples were mounted in ProLong Gold Antifade with DAPI (Thermo Fisher). For phalloidin staining, the basal 0.5-2.5 cm of leaf 4 from *o1* and wild-type siblings was fixed and stained with Alexa fluor 488-phalloidin (Thermo Fisher) as described previously (Cartwright et al., 2009; Nan et al., 2019). Nuclei and cell walls were stained using 10 µg mL−1 propidium iodide (Thermo Fisher).

### CFP-TUBULIN, YFP-TUBULIN, and TAN1-YFP imaging

Time-lapse imaging was performed by taking a Z-stack every 6 min and assessing the morphology of the mitotic structure. The start of metaphase was counted from the first time the spindle was observed until the anaphase spindle was observed. This time-point became the first time-point for anaphase. Telophase timing was measured from the first time-point a phragmoplast was observed until the phragmoplast was completely disassembled. Leaf No. 5 or 6 from 12 to 14 day-old maize seedlings were used. Samples were prepared as described before (19).

### *Dcd2-O* mapping

*dcd2-O* was mapped to chromosome 4 using a near isogenic line analysis after four backcrosses to B73. *dcd2* mutants in the B73 background were crossed to Mo17 and W64 inbred lines creating mapping populations for positional cloning. Markers on chromosome 4 were evaluated in over 1300 *dcd2* mutants from the two mapping populations. Markers used for fine mapping on chromosome 4 included SSR markers from Sigma’s Maize SSR Polymorphic Primer Set (umc2038, umc1620, bnlg1189, umc1871, and bnlg2162) and discovered SNP markers (N19-SNP – amplify with 5’cggagagaaaggtttggttg and 5’ctcatcgttccgtttggttt and cut with MboII, F07-SNP – amplify with 5’tggaataaacccagctttgc and 5’gccaaccagatgctcttctc and cut with StuI, and AC185621 – amplify with 5’aagtcaacctgttgcgttcc and 5’cgccttctgattcaccatct and cut with PvuII).

### Co-IP/MS

Co-IP/MS experiments were performed as previously described with some modifications (Facette et al., 2015). Families segregating *o1-N1242A* and wild-type siblings were used, with three biological replicates per genotype. The experiment was duplicated twice, using either the O1-11759 antibody or the o1-117560 antibody. The cell division zone (0.5 cm to 2.5 cm from the leaf base) were isolated from unexpanded leaves 4-6 of 10-to-14 day-old plants. Four to ten plants were pooled per replicate to obtain 1.5 grams of tissue. The leaf tissues were ground in liquid nitrogen. Extraction buffer (50mM Tris, pH 7.5, 150 mM NaCl, 5mM EGTA, 5mM EDTA, 0.3%-mercaptoethanol, 1% Plant Protease Inhibitor Cocktail) was added at 1 ml per 0.25 g of tissue and the mixture was homogenized for 3 × 15 seconds, with 30s breaks in between. Extracts were centrifuged at 15000 rpm in microcentrifuge two times, then the supernatant was transferred and centrifuged at 110,000xg for 45 min in ultracentrifuge. After spinning, the supernatant was removed and the pellet was resuspended in 500 µl (per 0.25g of starting tissue) of solubilization buffer (50 mM Tris pH 7.5, 150 mM NaCl, 1% NP-40, 10% glycerol, 0.05% Sodium deoxycholate). Samples were sonicated 2×15 seconds on ice and left rotating at 4°C for at 1-2 h. The extracts were centrifuged again at 110,000xg for 45 min, and the supernatant was transferred to a new tube. Dynabeads coupled with anti-O1 antibody 11759 and 117560) were prepared according to Dynabeads kit (Thermo Fisher) and added to the supernatant. The sample was incubated rotating at room temperature for 30 min. The Dynabeads-Co-IP complex was washed according to Dynabeads kit instructions.

The Dynabeads-Co-IP complexes were digested overnight at 37°C in 400ng of trypsin (Promega) per sample in 50 mM NH_4_CO_3_ buffer. After digestion, peptides were reduced with 1 mM dithiothreitol at room temperature for 30 min and then alkylated with 5 mM iodoacetamide at room temperature in the dark for 30 min. Formic acid was added to a 0.1% final concentration and peptides were extracted from the beads and desalted using the C18-Stage-Tip method and then vacuum dried. The dried peptides were reconstituted in 20µl of 5% formic acid/5% acetonitrile and 3 μl of sample was injected on LC column with 60 min. of gradient method for each run for MS analysis. Samples were run in technical triplicates on a Q-Exactive mass spectrometer with instrument and chromatography settings as described previously (Markmiller et al., 2018). The RAW files were analyzed using Andromeda/MaxQuant (version 1.6.0.16) (Cox and Mann, 2008) with the default settings except the match between the runs (MBR) and label free quantification (LFQ) settings were enabled. Data were searched against a concatenated target-decoy database comprised of forward and reverse FASTA peptide sequences from the B73 RefGen_v4, AGPv4 (Jiao et al., 2017).

## Acknowledgements

We would like to thank Miguel Vasquez, Lindy Allsman, Laurie Smith and Anne Sylvester for performing maize crosses. Research reported in this publication was supported in part by: NSF-IOS 1754665 to MRF, NSF-MCB 1716972 to CGR and NSF-CAREER-MCB 1942734 to CGR, and NIH NIGMS P30 GM110907. Microscope image data using the Zeiss LSM710 was gathered in the UNM CETI Cell Biology Core with support from NIH NIGMS P30 GM110907. Microscope image data using the Nikon A1R was gathered in the Light Microscopy Facility and Nikon Center of Excellence at the Institute for Applied Life Sciences, UMass Amherst with support from the Massachusetts Life Sciences.

## Author Contributions

QN, HL, JM, AJW, EJB, CGR and MRF designed the research. QN, HL, JM, LL, AF, and MRF performed the research; QN, HL, JM, LL, AF, EJB, CGR, and MRF analyzed the data; AW contributed the *dcd2* allele; MRF, CGR, HL, and QN wrote the paper with input from other authors; MRF, CGR, and EJB provided funding.

---

**Supplemental Figure 1.**
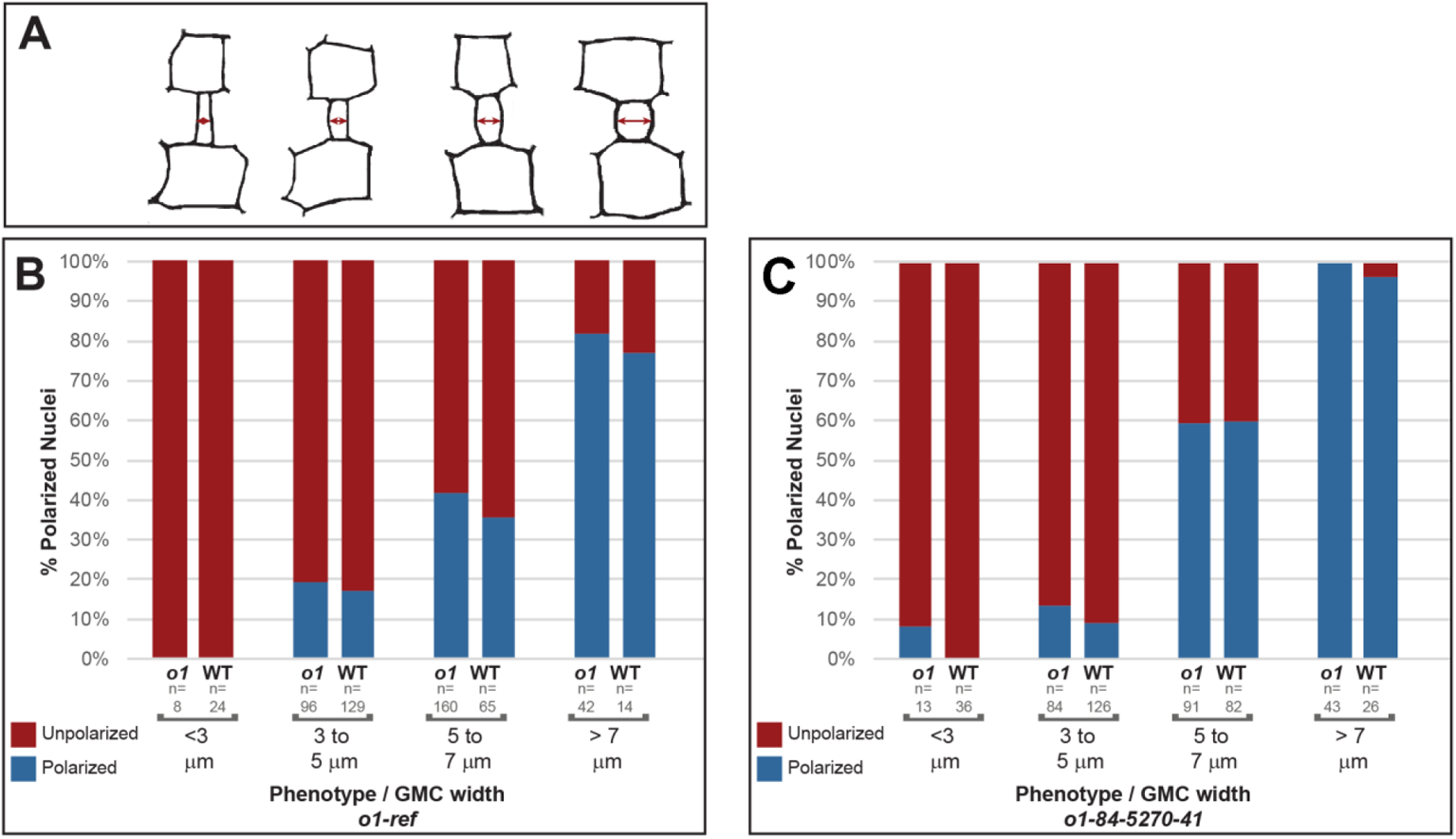
Nuclear migration is normal in *o1.* Additional alleles of *o1* were analyzed for nuclear polarization defects. Data from *o1-N1242A* are shown in the main text in Figure 4. (A) Cartoon depicting representative cell outlines at increasing GMC widths. Percent polarized nuclei in SMCs at progressive developmental stages in (B) *o1-ref* and (C) *o1-84-5170-40* mutants and their corresponding wild-type siblings. Developmental stage was inferred by the width of the adjacent GMC. Fisher’s exact tests comparing *o1* mutants and their respective wild-type siblings indicate no differences between mutants and wild type at each developmental stage (p>0.05 in all cases, with no testing correction).

**Supplemental Figure 2.**
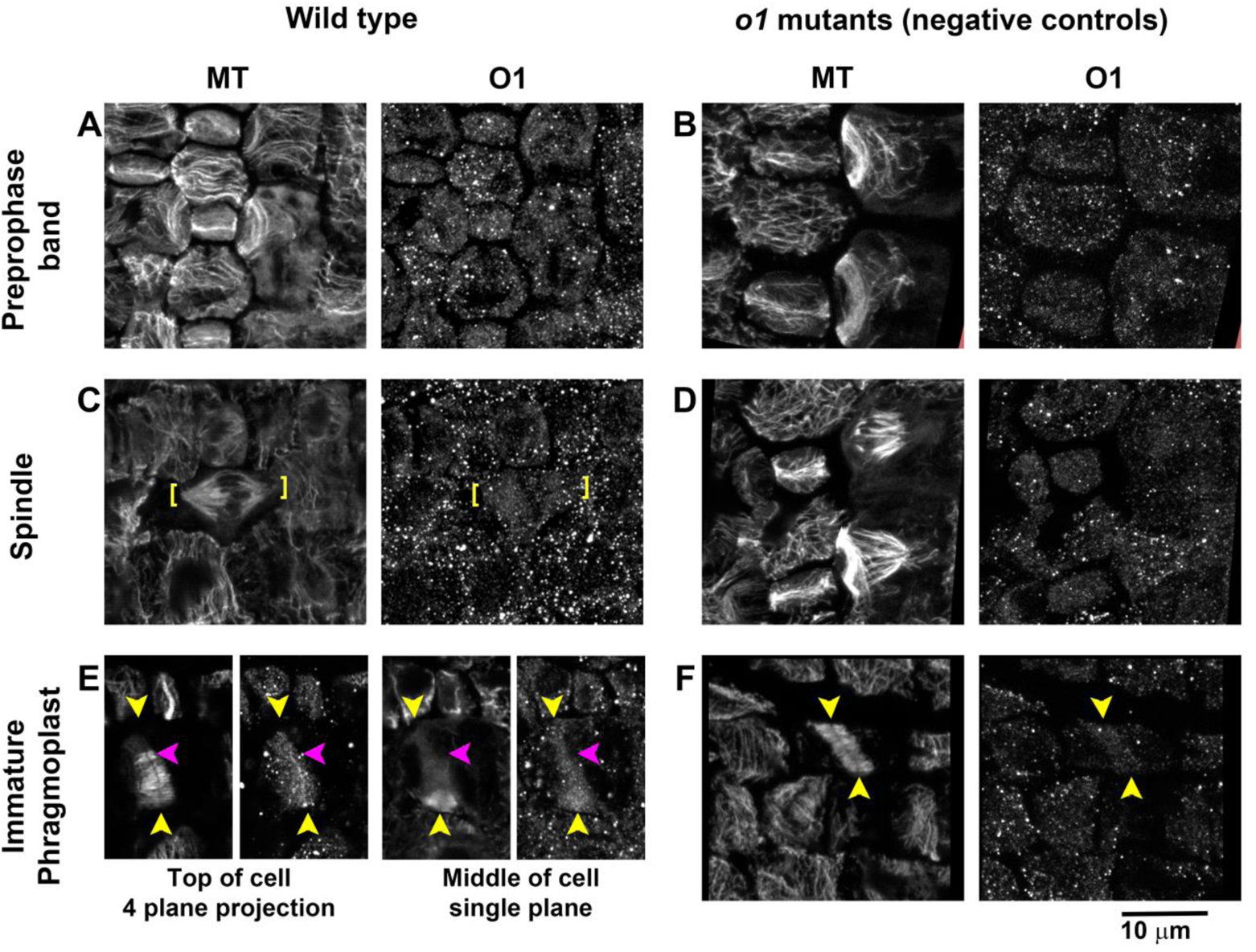
O1 localizes to phragmoplasts. Wildtype siblings (A,C,E) and *o1* mutants (B, D, F) were co-immunostained using an affinity-purified anti-O1 antibody and an anti-tubulin antibody. No specific accumulation of O1 was seen in preprophase bands (A,B) or spindles (C,D). Panel E shows a partially expanded phragmoplast in a wildtype cell. A bright line of O1 staining is seen at the phragmoplast midplane, which stops at the edge of the phragmoplast, indicating that O1 staining is not present at the cell cortex. Images in A, B and F are Z-projections of 4 planes. Images in C and D are Z-projections of 6 planes. All images at the same magnification.

**Supplemental Figure 3.**
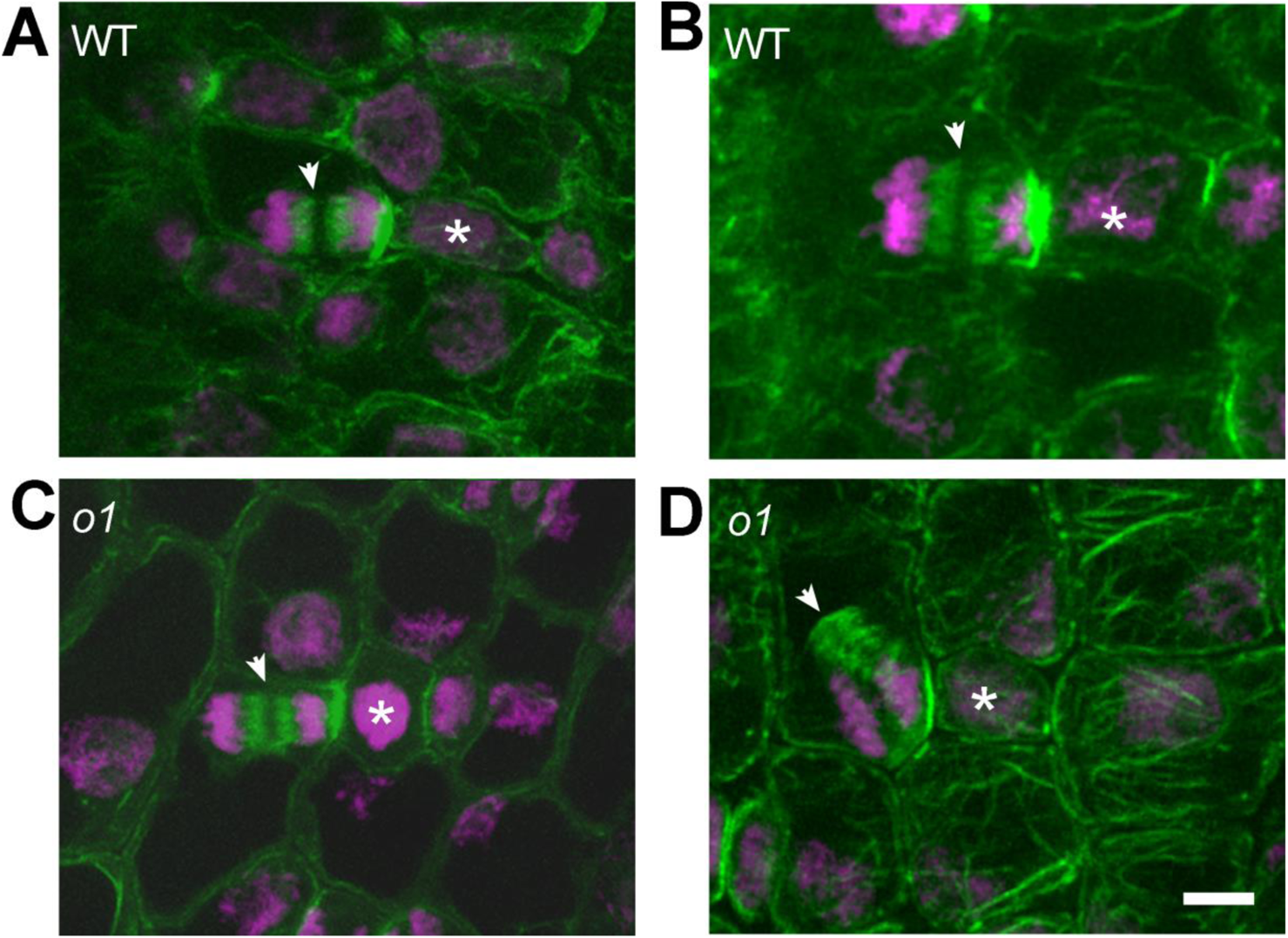
Actin in phragmoplasts in wildtype and *o1* mutants. Z-projections of Alexafluor488-phalloidin stained actin (green) and DAPI-stained DNA (magenta) in SMCs from wild type siblings (A, B) or *o1-N1242A* mutants (C, D). Arrowheads point to midline of phragmoplasts. Similar defects in phragmoplast guidance were observed when observing either microtubules or actin; only late stage phragmoplasts differed from wildtype and could become misguided or twisted, but typically had a clear midline. Scale bar = 5 micrometers.

**Supplemental Figure 4.**
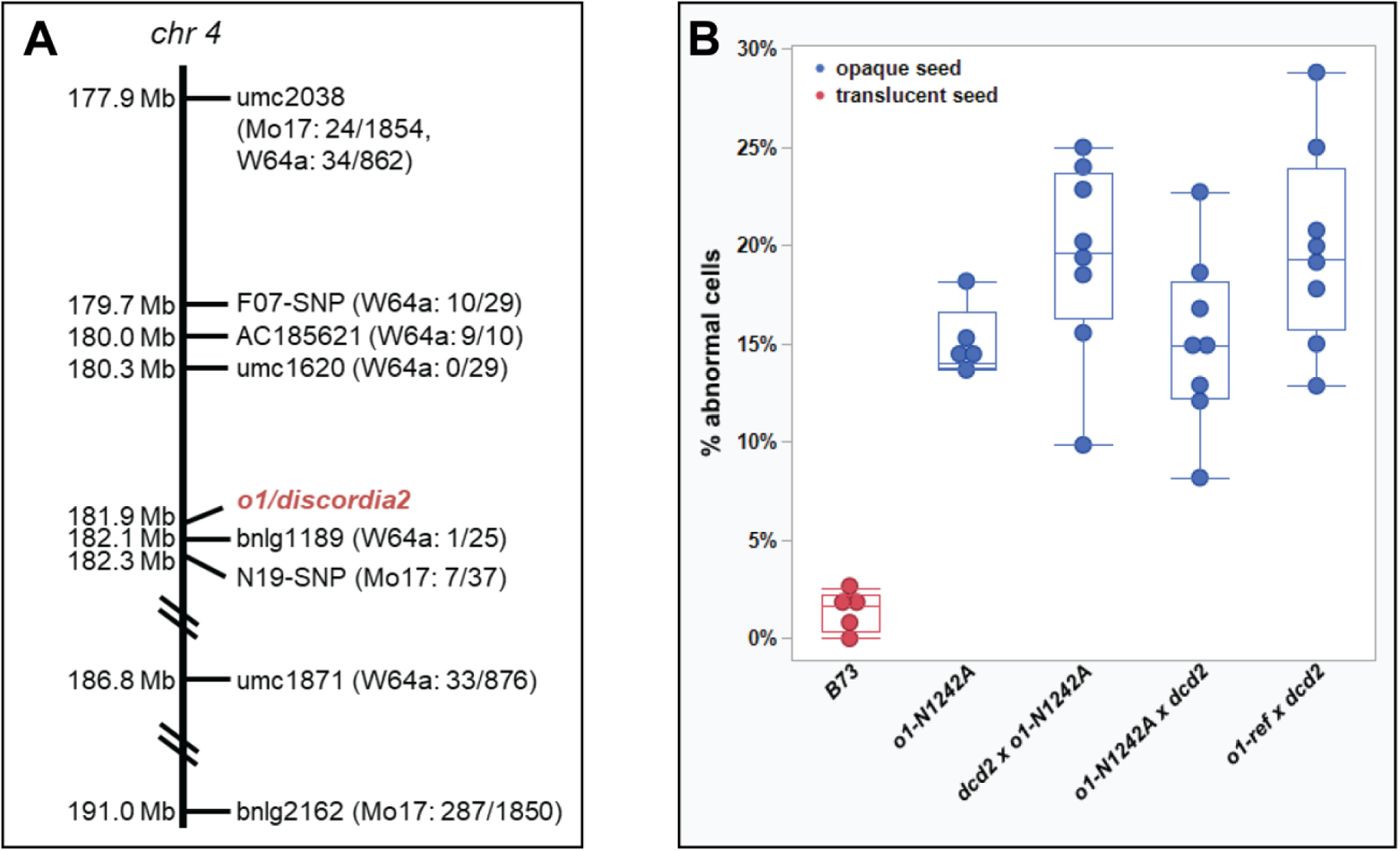
*Dcd2* is allelic to *O1.* (A) *dcd2-O* was mapped to chromosome 4 using a near isogenic line analysis after four backcrosses to B73. The line figure shows the positions of the markers on chromosome 4 and the number of recombinant chromosomes detected / total chromosomes screened for each marker. The position of *o1/dcd2* is also indicated. (B) Mutant *o1-N1242A* or *o1-ref* plants were crossed with *dcd2* plants and the F1 progeny was scored for opaque seeds. The cross to *o1-N1242A* was done with *dcd2* as both a female and as a male. For each cross, all seeds (hundreds per cross) were phenotypically opaque. Leaf 4 of seedlings was assayed for abnormal subsidiary cells. Between 50 and 150 cells were counted per plant. F1 progeny for all crosses had abnormal subsidiary cells at a frequency comparable to the *o1-N1242A* mutant allele grown at the same time, indicating *O1* and *Dcd2* are allelic.

**Supplemental Figure 5.**
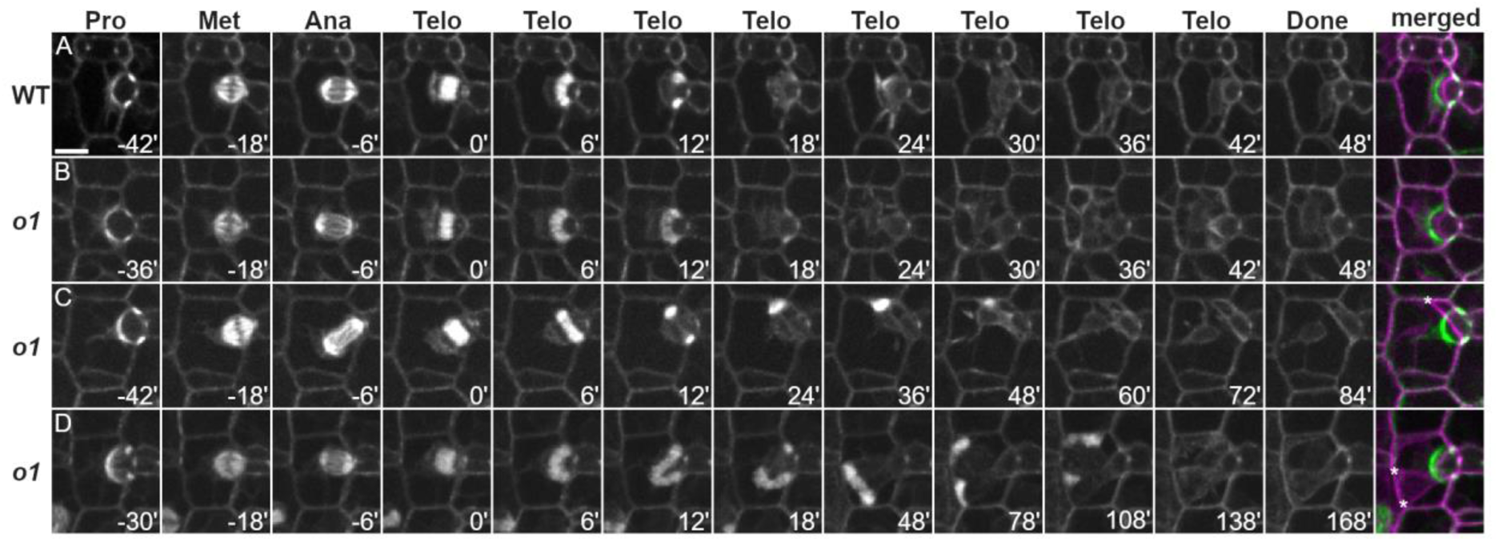
Time-lapse imaging of wild-type and *o1-84-5270-40* mutant subsidiary mother cell division. Progression of cell division was observed in dividing wild type and *o1-84-5270-40* SMCs from leaf 5 or 6 expressing CFP-TUB. (A) Wild-type cell division. In wild-type siblings, subsidiary mother cells all showed normal division plane orientation and the newly formed cell wall matched the former location of the PPB (n =78). (B) Correctly oriented *o1-84-5270-40* cell division. Approximately two thirds (n = 63/96) of phragmoplasts showed normal phragmoplast expansion. (C,D) Misoriented *o1-84-5270-40* cell divisions. Approximately one-third of *o1-5270-84* SMCs (33/96 = 34%) displayed misguided phragmoplasts, resulting in abnormal division planes. Pro - prophase; Met - metaphase; Ana - anaphase; Telo - telophase; Done - completed division; Merged - overlay of prophase (green) and completed division. Time (minutes) listed at the bottom of each image. Z-projections of 6 images. Misplaced cell walls are indicated by asterisks. All cells displayed at the same magnification; scale bar in A= 10 µm.

**Supplemental Figure 6.**
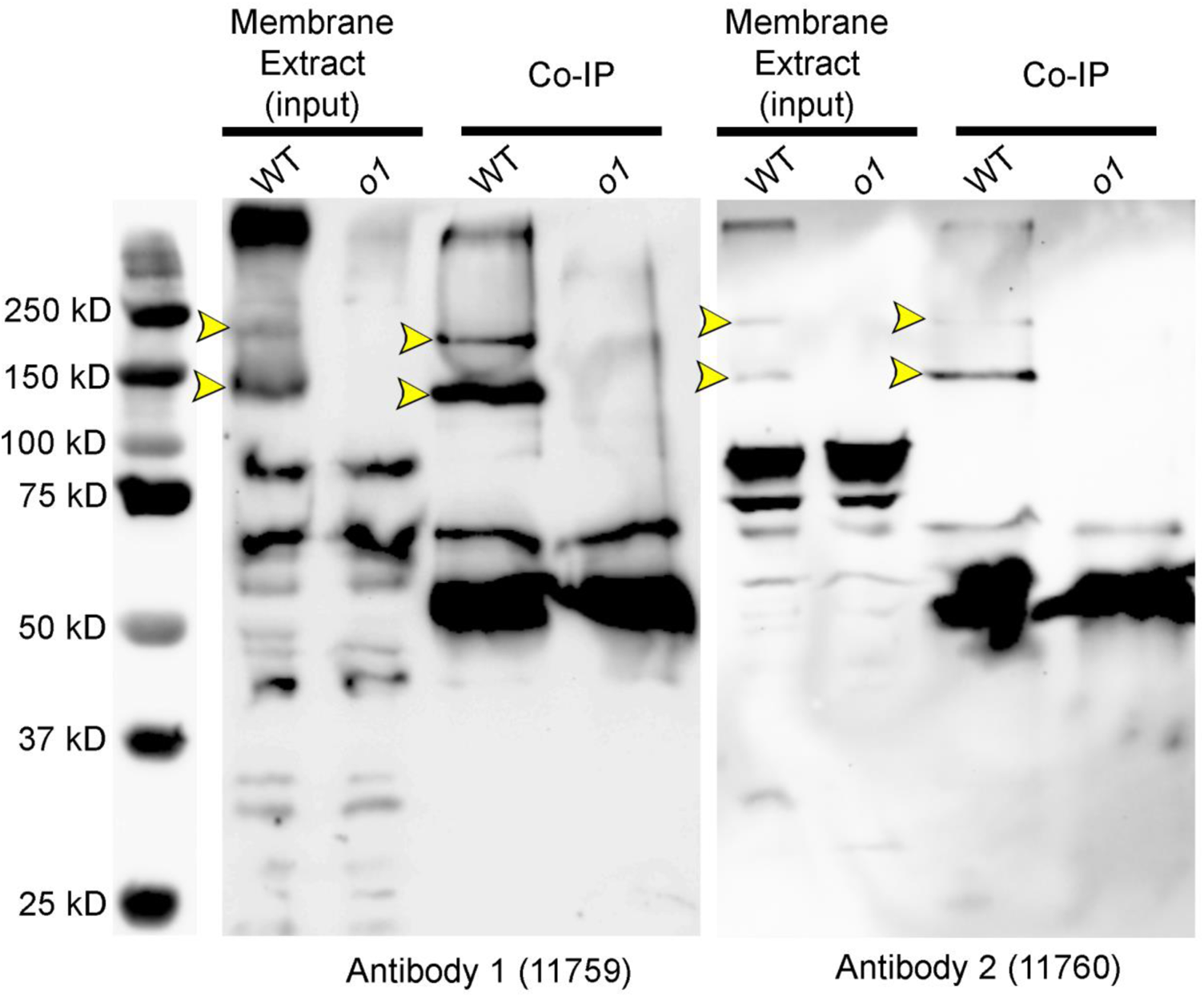
Validation of O1 antibodies and co-IP/western blot results. Membrane protein extracts from *o1-N1242A* mutants and wild type siblings were prepared, and co-IPs were performed. Immunodetection of proteins via western blot indicated that both antibodies detect two bands specific to WT that were missing in *o1* mutants (yellow arrowheads). O1 has multiple predicted splice isoforms; our western blot and co-IP/MS data are consistent with at least two active isoforms in dividing leaf tissue.

**Supplemental Figure 7.**
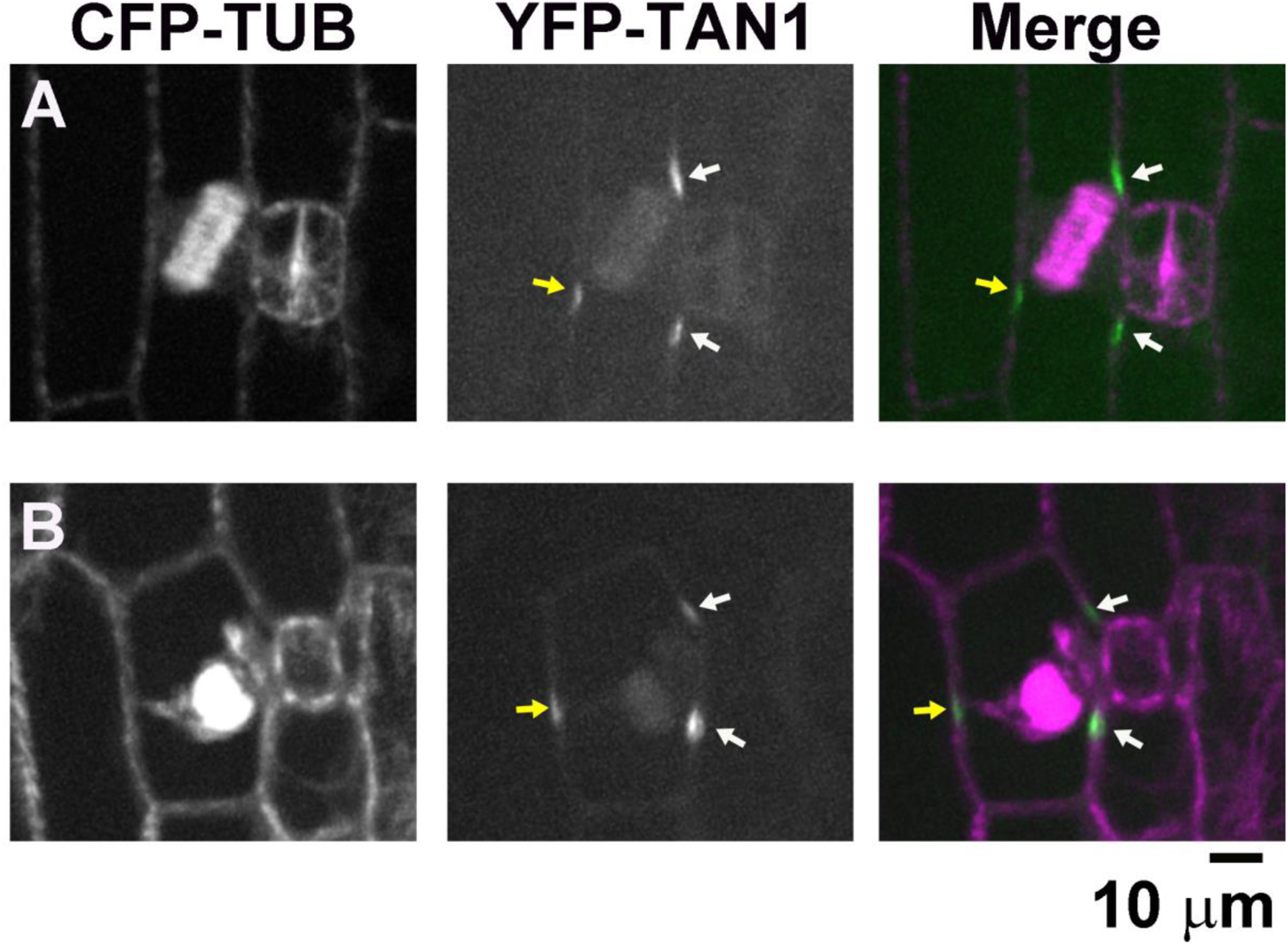
Unexpected TAN1-YFP localization in *o1* telophase SMCs. Dividing SMCs, from o1-N1242A leaf 5 or 6, co-expressing CFP-TUB and TAN-YFP were examined. Rarely (n=2/231), TAN1-YFP was observed not only at the expected cortical division site (white arrows), but also distal to the predicted division site(yellow arrows). In both cases, the phragmoplast had not yet met the cell cortex.

**Supplemental Table 1.** PAN1-YFP is polarized in *o1* subsidiary mother cells that have divided normally and abnormally. PAN1-YFP polarization was assayed in recently divided subsidiary cells from leaf 4 of 5 o1 plants and 4 wildtype siblings. Only subsidiary cells adjacent to normally formed guard mother cells were counted.

|  | Normal SMC division |  | Abnormal SMC Division |  |  |
| --- | --- | --- | --- | --- | --- |
|  | Polarized | Not Polarized | Polarized | Not Polarized | TOTAL |
| **o1** | 136 (65%) | 0 | 73 (35%) | 0 | 209 (100%) |
| **WT** | 294 (98%) | 0 | 5 (2%) | 0 | 299 (100%) |

**Supplemental Table 2.** Total number of SMCs examined for TAN1-YFP localization at different steps of the cell cycle. In all cells, TAN-YFP was located to the predicted division site. In 2 early telophase o1 cells, TAN1-YFP was located to the predicted division site in addition to another location.

|  | Stage of Cell Cycle |  |  |  |  |  |  |
| --- | --- | --- | --- | --- | --- | --- | --- |
| Genotype | Prophase | Metaphase | Anaphase | Early telophase | Late telophase |  | Total Cells |
|  |  |  |  |  | Normal | Misguided |  |
| **Wildtype** | 66 | 30 | 12 | 21 | 58 | 0 | **154** |
| **o1** | 85 | 19 | 13 | 32 | 38 | 44 | **231** |

**Supplemental Dataset 1**. Peptides identified in co-IP/MS experiments in *o1* mutants and wildtype siblings using two different antibodies.

**Movie S1**. **Normal asymmetric division of a wild-type SMC**. An example of wild-type SMC dividing using CFP-TUBULIN. The division begins in preprophase and proceeds until the end of telophase. The location of the new cell wall is accurately aligned with the previous preprophase band position. The video is playing at 5 frames per second, with each frame representing 6 min. Z projection of 3 images.

**Movie S2. Division of an *o1-N1242A* maize leaf subsidiary mother cell with a misguided phragmoplast.** An example of *o1* mutant subsidiary cell dividing using CFP-TUBULIN demonstrating a phragmoplast that becomes misguided shortly after meeting the cell cortex. The division begins in preprophase and proceeds until the end of telophase. The location of the new cell wall is not accurately predicted by the previous position of the preprophase band. The video is playing at 5 frames per second, with each frame representing 6 min.

**Movie S3. Division of an *o1-N1242A* maize leaf subsidiary mother cell with a misguided phragmoplast.** An example of *o1* mutant subsidiary cell dividing using CFP-TUBULIN demonstrating a phragmoplast that becomes misguided near the end of telophase. The division begins in preprophase and proceeds until the end of telophase. In this case, only a small divergence from the previously established division plane is observed. The video is playing at 5 frames per second, with each frame representing 6 min.

**Movie S4. Localization of TAN1-YFP in *o1-N1242A* SMC with phragmoplast** 3-D projection of *o1-N1242A* expressing TAN1-YFP (yellow) and CFP-TUB (cyan). The two lower SMCs have preprophase bands marked by CFP-TUB; only one cell has thus far accumulated TAN1-YFP at the cortical division site. The top SMC shows TAN1-YFP correctly marking the cortical division site, but the phragmoplast has become misguided. A correctly formed subsidiary cell and sister pavement cell (showing nucleolar TAN1-YFP localization) are also shown at the top.

